# WDR5 promotes breast cancer growth and metastasis via KMT2-independent translation regulation

**DOI:** 10.1101/2022.03.30.486357

**Authors:** Wesley L Cai, Jocelyn F Chen, Huacui Chen, Emily Wingrove, Sarah J Kurley, Lok Hei Chan, Meiling Zhang, Anna Arnal-Estapé, Minghui Zhao, Amer Balabaki, Wenxue Li, Xufen Yu, Yali Dou, Yansheng Liu, Jian Jin, Thomas F Westbrook, Don Nguyen, Qin Yan

## Abstract

Metastatic breast cancer remains a major cause of cancer related deaths in women and there are few effective therapies against this advanced disease. Emerging evidence suggests that key steps of tumor progression and metastasis are controlled by reversible epigenetic mechanisms. Using an *in vivo* genetic screen, we identified WDR5 as an actionable epigenetic regulator that is required for metastatic progression in models of triple-negative breast cancer. We found that knockdown of WDR5 in breast cancer cells independently impaired their tumorigenic as well as metastatic capabilities. Mechanistically, WDR5 promotes cell growth by increasing ribosomal gene expression and translation efficiency in a KMT2-independent manner. Consistently, pharmacological inhibition or degradation of WDR5 impedes cellular translation rate and the clonogenic ability of breast cancer cells. Furthermore, combination of WDR5-targeting with mTOR inhibitors leads to potent suppression of translation and proliferation of breast cancer cells. These results reveal novel therapeutic strategies to treat metastatic breast cancer.

## Introduction

In the United States, metastatic breast cancer is the second leading cause of cancer related death among women (Harbeck et al., 2019; Torre et al., 2017). In particular, triple-negative breast cancer (TNBC) has the worst prognosis among all breast cancer subtypes, largely owing to its high metastatic proclivity to the lungs and other sites and few effective treatments against this disease once it has metastasized (Al-Mahmood et al., 2018). Recently developed targeted therapies for TNBC, including poly (ADP-ribose) polymerase (PARP) inhibitors or immune checkpoint inhibitors, are effective in patients whose tumors express BRCA1/2 mutations or high programmed death-ligand 1 (PD-L1), respectively (Gonzalez-Angulo et al., 2011; Lyons & Traina, 2019). However, these patients account for only 9.3-15.4% of TNBC cases (Armstrong et al., 2019) and new treatment strategies are urgently needed.

Emerging evidence suggests tumor growth is modulated by reversible epigenetic mechanisms (Blair & Yan, 2012; Cao et al., 2014; Cao & Yan, 2013; Chen & Yan, 2021). In primary human breast cancers, we and others recently identified distinct chromatin states, which distinguish established molecular subtypes and correlates with metastatic relapse and poor clinical outcome (Cai et al., 2020). Therefore, regulators of histone modifications and chromatin dynamics in particular, may be required for breast cancer progression. The identity of such regulators as well as strategies to therapeutically target them in the metastatic setting remain unclear.

Epigenetic regulators that are known to be involved in tumorigenesis include the KMT2 (also known as MLL/SET1) family protein complexes which marks active promoters and enhancers with H3K4 methylation, and the non-specific lethal (NSL) complex which acetylates histones (Dias et al., 2014; Raja et al., 2010; Ruthenburg et al., 2007; Wysocka et al., 2005). WDR5 is a WD40 repeat protein that scaffolds the assembly of the KMT2 and NSL complexes (Guarnaccia & Tansey, 2018). More recently, WDR5 was found to physically interact with the proto-oncogene and transcription factor MYC to guide its chromatin binding and transcriptional activation, suggesting that WDR5 is a tractable target for MYC-driven cancers (Thomas, Foshage, et al., 2015; Thomas, Wang, et al., 2015). Aberrant WDR5 expression itself may occur in a number of cancer types (Chen et al., 2015; Dai et al., 2015; Ge et al., 2016). Biologically, WDR5 may contribute to tumor sphere formation and cell proliferation (Carugo et al., 2016; Chung et al., 2016). Molecularly, WDR5 is reported to modulate the expression of various genes which may be specific to cell type or cell state (Bryan et al., 2020; Oh et al., 2020). On the other hand, WDR5 was recently discovered to broadly regulate the expression of ribosomal protein (RP) genes across multiple cell lines and cancer types (Aho et al., 2019; Bryan et al., 2020; Guarnaccia et al., 2021). Moreover, deregulation of RP gene expression and translation have been implicated in breast cancer metastasis (Ebright et al., 2020). However, the relative importance of different WDR5 effector functions and their requirement for breast cancer progression and metastasis have not been well-studied.

Here, we establish an *in vivo* screening platform that identified WDR5 as a key regulator of breast cancer cell growth and metastatic colonization. We further show that WDR5 regulates RP gene expression and global protein translation independently of the KMT2 complex. Moreover, our results indicate that WDR5 inhibition or degradation could be used as a therapeutic approach for TNBC, and that WDR5 targeting could be combined with mTOR inhibitors to achieve significant therapeutic benefit.

## Results

### The establishment of *in vivo* lung metastasis screening platform

To identify actionable epigenetic targets for breast cancer metastasis, we conducted parallel *in vivo* and *in vitro* functional screens using an inducible, barcoded shRNA library (**Figure 1A**). We first compiled a list of epigenetic regulators based on: 1) if they could be targeted with existing pharmacological agents, 2) if their expression correlated with poor survival in multiple independent datasets (hazard ratio>1 and p-value<0.05) or, 3) if their expression was increased in the lung metastatic cell sub-population MDA-MB-231-LM2 (LM2) cells when compared to the parental TNBC cell line MDA-MB-231. We designed our screen using the LM2 cells because they reproducibly generate lung metastasis and lung is the most frequent site of distant relapse in TNBC patients (Lin et al., 2008; Minn et al., 2005). Accordingly, we tested the knockdown efficiency of 336 shRNAs targeting 100 epigenetic regulators and selected one shRNA with the best knockdown efficiency per target gene. We then subcloned these shRNAs into the doxycycline (DOX) inducible and barcoded pINDUCER10 lentivirus to generate a focused knockdown-validated shRNA library (**Figure 1-figure supplement 1A**) (Meerbrey et al., 2011). Included in this library were positive control shRNAs against *BUD31 (*shBUD31) and *SAE2* (shSAE2), which were previously shown to be essential for LM2 cell proliferation, along with shRNAs against *CHEK1* and *STAMBP,* which served as negative controls (Hsu et al., 2015; Kessler et al., 2012).

**Figure 1.**
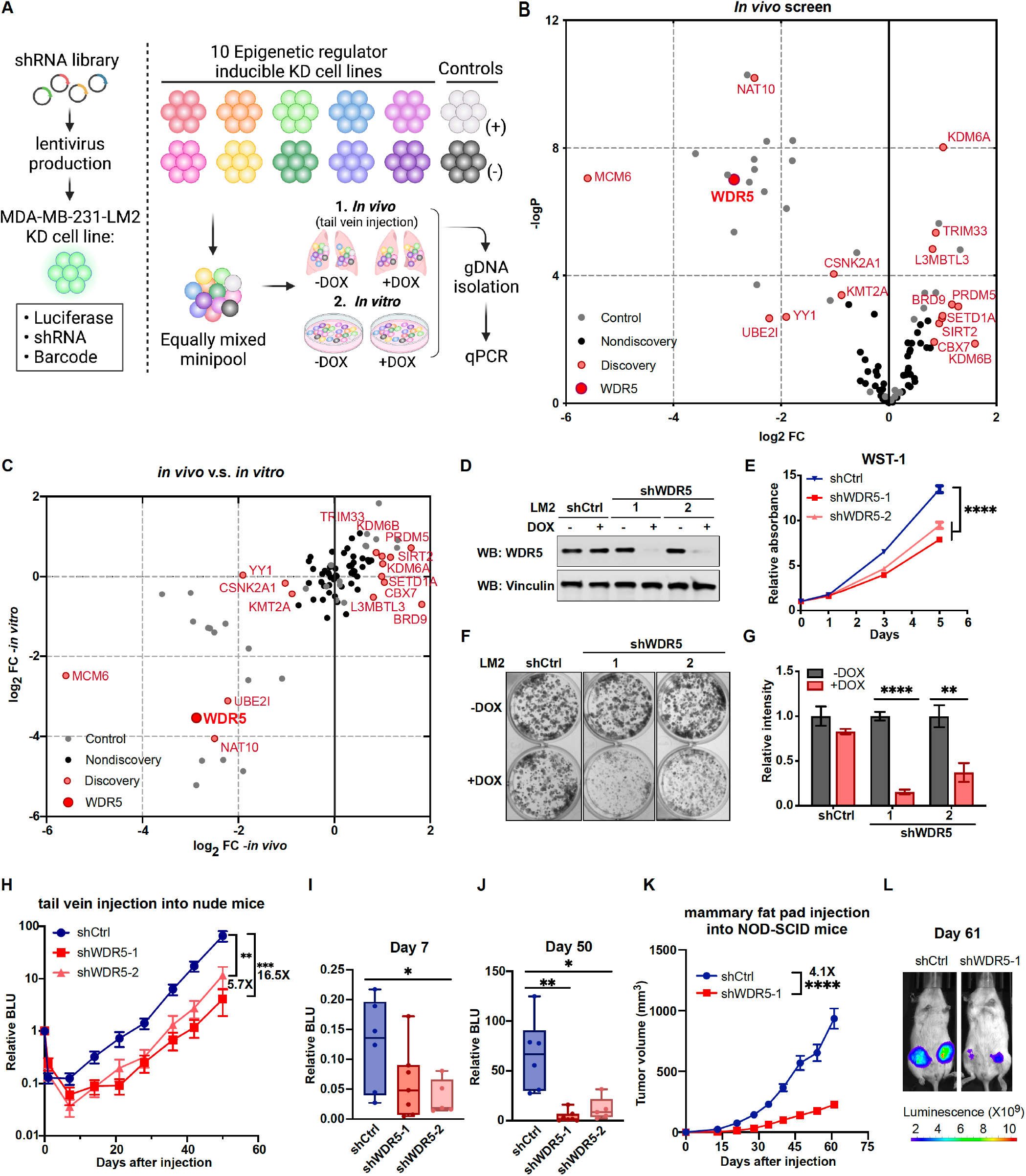
Decreasing WDR5 reduces breast cancer cell growth and lung metastasis. (A) Schematic of *in vivo* and *in vitro* screening work flow. Epigenetic regulator inducible knockdown cell lines were equally mixed and injected into mice intravenously or cultured under control or doxycycline (1 μg/mL) treated condition. Both lungs (*in vivo*) and cells (*in vitro*) were harvested for gDNA and subjected to barcode qPCR as the screening output. (B) Volcano plot showing the results of the *in vivo* screen. Each data point is an average of 10-20 mice. Discovery hits are selected using P<0.05 and log_2_FC (Fold Change) >0.8 or log_2_FC<-0.8. (C) Log_2_FC of the *in vivo* screen results versus log_2_FC of the *in vitro* screen results for each of the epigenetic regulators. Discovery hits are selected as in (B). (D) Western blot analysis of the indicated proteins in LM2 cells harboring inducible control or WDR5 targeting (shWDR5-1 and shWDR5-2) shRNA after 3 days of doxycycline (1 μg/mL) induction. (E) WST-1 proliferation assays of LM2 cells from (D) after indicated days of doxycycline (1 μg/mL) treatment. Each symbol indicates mean ± SD for representative experiment performed in quadruplicate (n=4, unpaired two-side Student’s *t* test). (F-G) Colony formation assays of LM2 cells from (D) after 9 days of either control or doxycycline (1 μg/mL) treatment. Representative images (F) and quantification (G) are shown (n=3, unpaired two-side Student’s *t* test). (H) Normalized bioluminescence signals of lung metastasis of mice injected intravenously with LM2 cells from (D) and kept under doxycycline chow. The data represent mean ± SEM (shCtrl: n=6; shWDR5-1: n=7; shWDR5-2: n=5). (I-J) Box plots of relative bioluminescence of indicated cell line at day 7 (I) and day 50 (J) post injection normalized to its day 0 value. (K) Tumor volume measurements of mice injected into the 4^th^ mammary fat pad with LM2 cells harboring inducible control or shWDR5-1. The data represent mean ± SEM. (L) Representative bioluminescence images of mice in (K) at day 61. Significance determined using unpaired two-tailed Mann-Whitney test (shCtrl: n=14; shWDR5: n=13). *p<0.05; **p<0.01; ***p<0.001; ****p<0.0001. For gel source data, see Figure 1-source data1.

LM2 cells were infected with individual shRNAs from this library and then all resulting cell lines were pooled together in equal numbers and cultured in either control or DOX conditions for up to 10 doublings. We then extracted genomic DNA (gDNA) from the pooled shRNA infected cells collected. qPCR analysis of gDNA confirmed that shBUD31 and shSAE2 were significantly depleted in the doxycycline-treated pools (**Figure 1-figure supplement 1B**). On the other hand, the amount of shSTAMBP and shCHEK1 expressing cells did not change significantly in either control or DOX conditions (**Figure 1-figure supplement 1C**). We next determined whether our controls perform similarly *in vivo*. Towards this end, LM2 cells expressing individual shRNAs from our library cells were combined into mini-pools, each containing 8-10 knockdown cell lines to ensure that any particular hairpin had enough representation and was above the detection limit *in vivo*. Mini-pools were then injected intravenously and treated with either control or DOX (in animal chow for *in vivo* conditions). After 50 days, tumor bearing lung tissue was collected and processed for gDNA extraction. We then compared the barcode abundance between control and DOX-treated lung tissue using qPCR analysis of gDNA. The results were normalized to the Day 0 value.

We found that DOX treatment does not in itself affect the *in vivo* lung metastatic growth kinetics (**Figure 1-figure supplement 1D**). In a representative minipool, shBUD31 and shSAE2 consistently dropped-out in the DOX-treated condition, whereas shSTAMBP remained unchanged (**Figure 1-figure supplement 1E**). shCHEK1 was enriched significantly, which is likely indirectly due to the depletion of other shRNA expressing cell lines in the mini-pools (**Figure 1-figure supplement 1E**). We screened a total of 69 genes in *in vivo* and *in vitro* conditions by splitting the entire shRNA library into 7 mini-pools. From the *in vivo* screen, we identified 16 significant hits (p<0.05, log_2_FC>0.8 or log_2_FC<-0.8, FC: +DOX/-DOX), and among these, 7 were drop-out hits where shRNA representation significantly decreased while 9 were enrichment hits where shRNA representation significantly increased (**Figure 1B**). Many of these *in vivo* drop-out candidates also showed drop-out phenotypes *in vitro* (**Figure 1C**). For example, our screen identified drop-out shRNAs against MCM6, an essential eukaryotic genome replication factor, and CSNK2A1, previously shown to enhance metastatic growth of MDA-MB-231 cells (Bae et al., 2016) (**Figure 1B, 1C**). Thus, many of these epigenetic targets may be, at least in part, required for the cell intrinsic fitness of metastatic cells.

### Decreasing WDR5 reduces breast cancer cell growth and lung metastasis

Among the top hits and potential therapeutic targets, we focused on WDR5, because it can be inhibited by small molecules and shWDR5 was the second-most significantly depleted shRNA *in vivo* (after shMCM6). WDR5 is known canonically as a scaffolding protein that recognizes and binds to methylated H3K4, allowing the modification of H3K4 tri-methylation by the KMT2 protein complex (Wysocka et al., 2005). More recently, WDR5 has been discovered to physically interact with and guide MYC to its transcriptional targets (Thomas, Wang, et al., 2015). WDR5 has also been implicated in the growth of metastatic breast cancer cells, although the mechanism underlying this function of WDR5 is not clearly delineated (Punzi et al., 2019).

We first confirmed the knockdown effect of WDR5 in individual un-pooled LM2 cells and by using independent shRNAs against WDR5 (shWDR5-1 and shWDR5-2). Following 3 days of shRNA induction *in vitro* (**Figure 1D**), both shRNAs against WDR5 caused a significant but modest decrease in cell proliferation when compared to a control shRNA (shCtrl) over 5 days (**Figure 1E**). In addition, using long-term *in vitro* colony formation assays over 9 days, we found a profound impact of WDR5 knockdown on the *in vitro* clonogenic ability of LM2 cells (**Figure 1F, 1G**). We next asked whether both shRNAs affect lung metastasis outgrowth *in vivo*. We induced knockdown for 3 days *in vitro* before injecting LM2 cells into the tail vein of mice and monitoring lung metastatic colonization and outgrowth over 50 days. We observed a significant impairment on lung colonization by LM2 cells as early as day 7 post-injection (**Figure 1H, 1I**). At the end point (day 50), the average lung metastatic burden in the mice with shWDR5 cells was 5.7 or 16.5-fold lower than that in mice with LM2 cells expressing the control shRNA (**Figure 1H, 1J**). In addition to metastatic colonization from circulation, we tested whether knockdown of WDR5 affects tumor growth and metastasis from the orthotopic mammary fat pad. We observed a significant decrease in mammary tumor growth in the shWDR5 group compared to control tumors (**Figure 1K, 1L, Figure 1-figure supplement 1F-H**). Notably, we observed an even larger decrease in lung and liver metastasis from the mammary fat pad tumors in the shWDR5 group as compared to shCtrl, which suggests the potential metastasis-specific function of WDR5 (**Figure 1-figure supplement 1 H-J**). Taken together, we showed that WDR5 is independently required for the cellular outgrowth, tumorigenic and lung colonizing capacities of LM2 TNBC cells.

### WDR5 depletion significantly reduces breast cancer cell growth across multiple breast cancer subtypes

Next, we tested the requirement for WDR5 in other cell line models and from distinct breast cancer subtypes. To this end, we silenced WDR5 with shWDR5-1 in additional breast cancer lines spanning three established molecular subtypes: TNBC (MDA-MB-453, HCC1143), estrogen receptor positive (ER+) (MCF7, T47D, MDA-MB-361), and HER2+ (UACC893, BT474, SKBR3) (**Figure 2A**). WDR5 silencing significantly reduced the clonogenic outgrowth of all the tested cell lines (**Figure 2B, 2C**), suggesting that WDR5 enhances tumor cell growth across multiple breast cancer subtypes. As we were particularly interested in evaluating the therapeutic potential of targeting WDR5 in TNBC, we first tested the efficacy of a known WDR5 inhibitor, OICR-9429, which is a small molecule antagonist of WDR5-KMT2 interaction (Grebien et al., 2015). We treated LM2 cells with OICR-9429 at 20 μM for 9 days and this also significantly reduced their colony formation ability (**Figure 2D, 2E**). Similar results were found when using OICR-9429 to treat two other TNBC cell lines, MDA-MB-453 and 4T1, although we noted that growth inhibition was more significant in MDA-MB-453 cells when using 30 μM of OICR-9429 (**Figure 2F, 2G, Figure 2-figure supplement 1A, 1B**).

**Figure 2.**
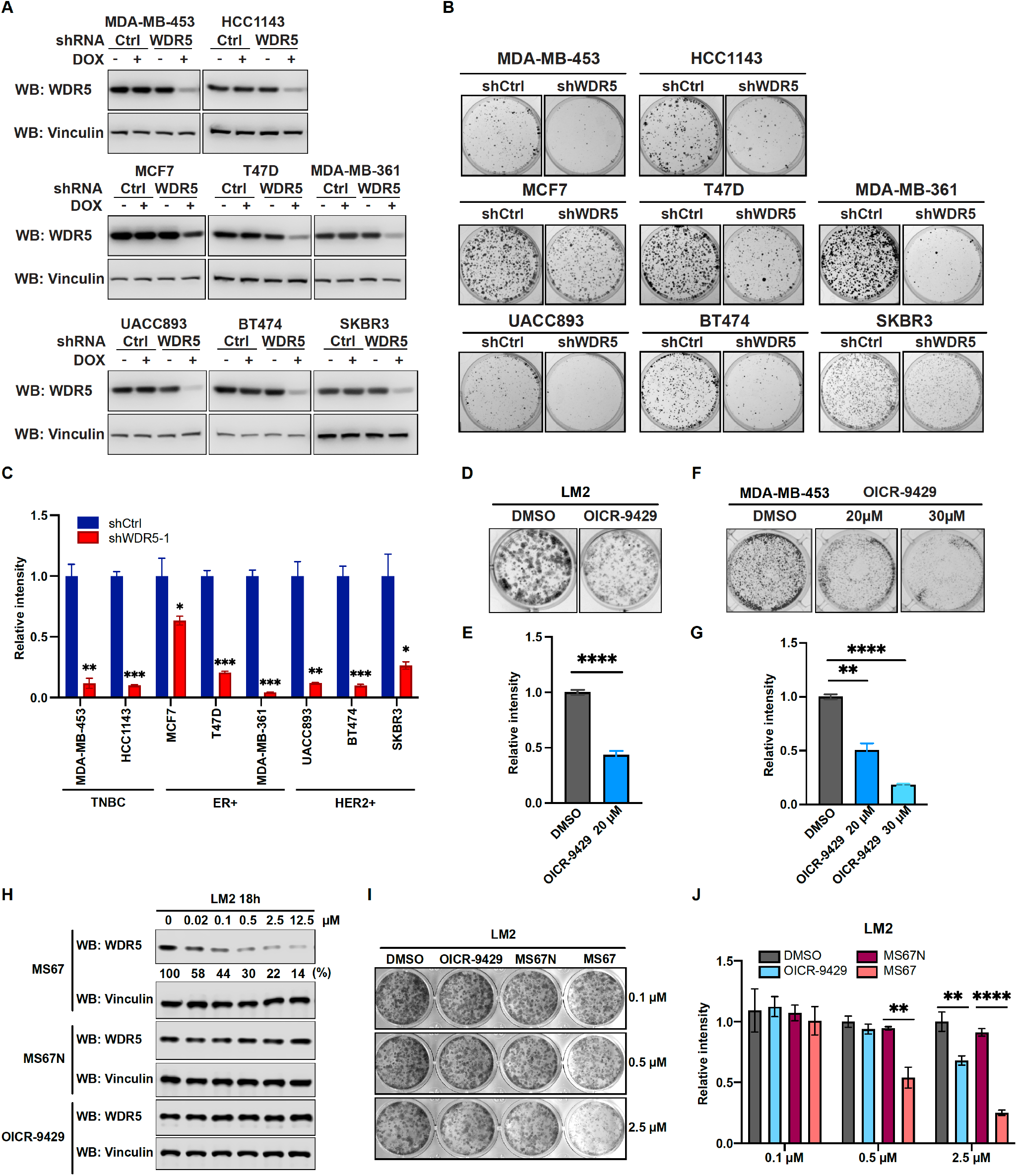
WDR5 targeting significantly reduces breast cancer cell growth across breast cancer subtypes. (A) Western blot analyses of WDR5 in the indicated cell lines infected with either control or WDR5-targeting hairpins with or without 3 days of doxycycline (1 μg/mL) induction. (B-C) Colony formation assays of indicated control or shWDR5-1 cell lines from (A) after 9 days of doxycycline (1 μg/mL) treatment. Representative images (B) and quantification (C) are shown (n=3, unpaired two-side Student’s *t* test). Cell lines are grouped by breast cancer molecular subtype. (D-G) Colony formation assays of LM2 (D) and MDA-MB-453 (F) after 9 days of either control or OICR-9429 treatment at indicated concentration. Representative images (D&F) and quantification (E&G) are shown (n=3, unpaired two-side Student’s *t* test). (H) Western blot analysis of WDR5 in LM2 treated with MS67, MS67N, or OICR-9429 at the indicated concentration for 18 hours. Band intensities of WDR5 were quantified by image J and normalized by those of vinculin control. (I-J) Colony formation assays of LM2 after 9 days of treatment with control, OICR-9429, MS67N, or MS67 at the indicated concentration. Representative images (I) and quantification (J) are shown (n=3, unpaired two-side Student’s *t* test). *p<0.05; **p<0.01; ***p<0.001; ****p<0.0001. For gel source data, see Figure 2- source data1-2.

As the effective concentration of OICR-9429 is relatively high and may lead to off-target effects, we sought to test the effect of our recently published WDR5 degrader MS67, which recruits WDR5 to Cullin4-CRBN E3 ubiquitin ligase complex for proteasome-mediated degradation (Yu et al., 2021). We first evaluated the effect of MS67 on degrading WDR5 in LM2 and MDA-MB-453 cells. We found that MS67, but not the negative control MS67N, which does not bind CRBN, nor OICR-9429, induced WDR5 degradation at a concentration as low as 0.02 μM (**Figure 2H, Figure 2-figure supplement 1C**). Specifically, at 2.5 μM MS67, we achieved ∼80% WDR5 degradation in LM2 cells and ∼70% of degradation in MDA-MB-453 cells (**Figure 2H, Figure 2-figure supplement 1C**). Additionally, the maximal degradation can be achieved at 8 hours post treatment and this effect remains stable for 72 hours in both LM2 and MDA-MB-453 cells (**Figure 2-figure supplement 1D, 1E**). Finally, we compared MS67-induced WDR5 degradation to OICR-9429 treatment on the clonogenic outgrowth of LM2 and MDA-MB-453 cells. We found that MS67 leads to ∼50% growth inhibition at 0.5 μM and ∼80% inhibition at 2.5 μM (**Figure 2I, 2J, Figure 2-figure supplement 1F, 1G**). Importantly, the effect of 2.5 μM MS67 treatment is comparable to shRNA knockdown and more potent than 20 μM OICR-9429 treatment in LM2 and MDA-MB-453 cells, while 2.5 μM of OICR-9429 treatment only caused a modest effect (**Figure 1F, 1G, 2B, 2C, 2I, 2J, Figure 2-figure supplement 1F, 1G)**. In summary, MS67-mediated WDR5 degradation showed improved growth inhibition of breast cancer cells when compared to the OICR-9429 compound.

### WDR5 targeting decreases ribosomal protein gene expression and global translation rates

To identify the molecular effects of WDR5 depletion in TNBC cells, we performed transcriptomic profiling of control and WDR5 knockdown cells. In addition to the MDA-MB-231 lung metastatic LM2 cells, we also tested the effect of WDR5 knockdown in independent MDA-MB-231 subpopulations that metastasize more readily to the brain (BrM3) or bone (BoM) (Bos et al., 2009; Kang et al., 2003). Because WDR5 has previously been shown to facilitate active transcription (Ang et al., 2011; Wysocka et al., 2005), we used spike-in RNA for normalization and found no changes in global RNA levels 3 days after shWDR5-1 induction (Jiang et al., 2011). Our analyses identified differentially expressed genes (DEGs) in all 3 organotropic-metastatic cell sub-populations (**Figure 3A**). In general, inhibition of WDR5 led to more down-regulated genes than up-regulated genes, which supports previous findings that WDR5 generally promotes transcriptional activation (Wysocka et al., 2005) (**Figure 3A)**. Certain DEGs were preferentially regulated in LM2, BrM3 or BoM cells, suggesting that WDR5 can regulate genes in a manner that is dependent on the metastatic proclivities of different breast cancer cell sub-populations (**Figure 3B**). On the other hand, we also found DEGs (264 down-regulated and 118 up-regulated) that were shared across all lines (**Figure 3B, Table S1**), indicative of some conserved WDR5 function in metastatic breast cancer cells.

**Figure 3.**
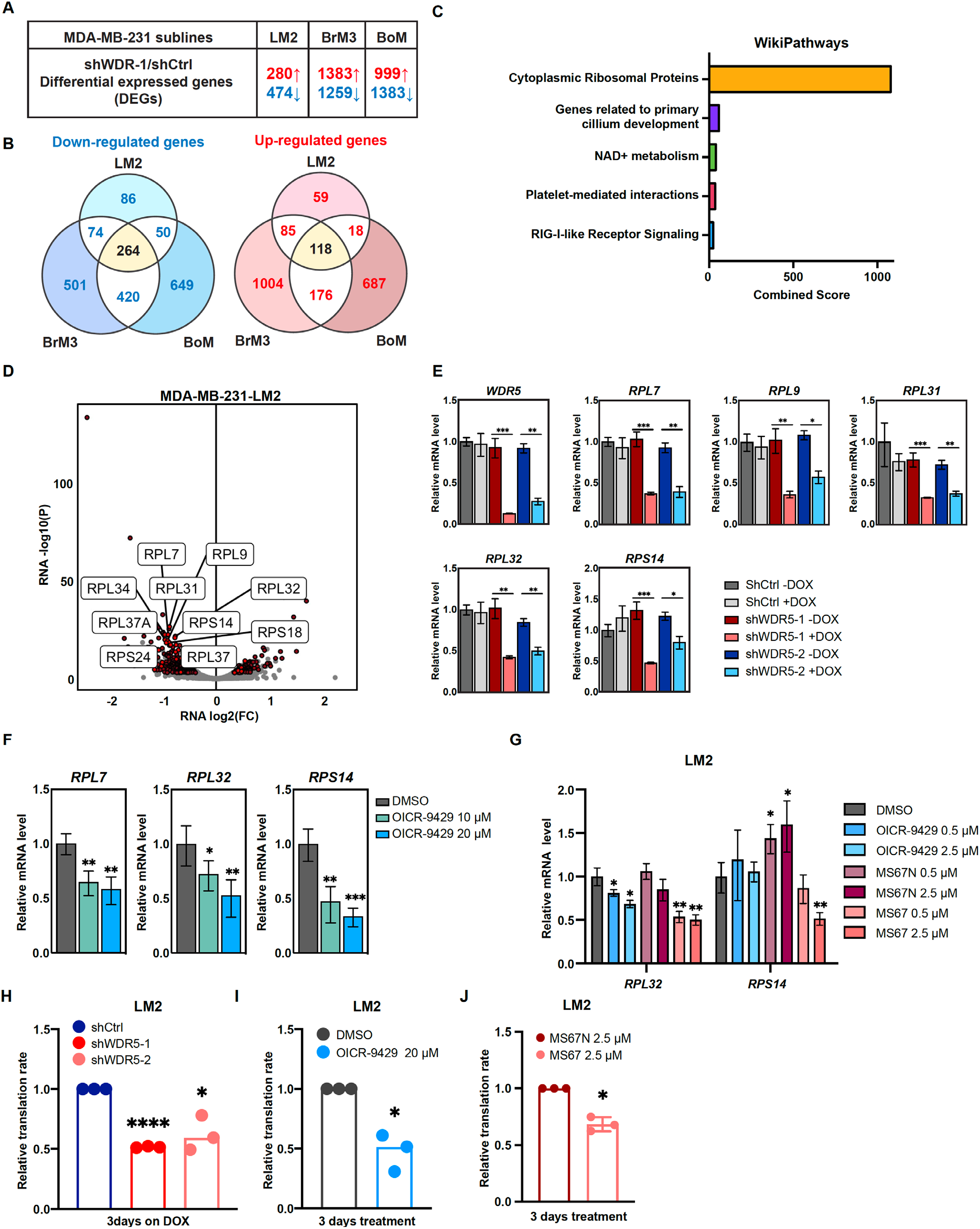
WDR5 targeting decreases ribosomal protein gene expression and global translation rates. (A) Table summarizing the number of differentially expressed genes (DEGs) after WDR5 silencing (red indicates up-regulated; blue indicates down-regulated) across three MDA-MB-231 organotropic sublines (LM2-lung; BrM3-brain; BoM-bone). (B) Venn diagram showing the number of overlap or distinct down-regulated genes (left) and up-regulated genes (right) after WDR5 silencing in the MDA-MB-231 organotropic sublines. (C) Gene ontology results using the down-regulated gene set shared by all three MDA-MB-231 organotropic sublines analyzed with Enrichr. (D) Volcano plot of DEGs after WDR5 knock-down in LM2. Shared DEGs across all lines highlighted in dark red and RPs (RPL and RPS) highlighted in light red. The top ten differentially expressed RPs are labelled. (E) RT-qPCR validation of selected DEGs in LM2 cells harboring shCtrl, shWDR5-1, or shWDR5-2 after doxycycline (1 μg/mL) induction for 3 days. (F) RT-qPCR validation of selected DEGs in LM2 cells after DMSO or OICR-9429 treatments at the indicated concentration for 3 days. (G) RT-qPCR validation of selected DEGs in LM2 cells after DMSO, OICR-9429, MS67N, or MS67 treatments at the indicated concentration for 48 hours. Significance determined by comparing each treatment to DMSO control (n=4, unpaired two-side Student’s *t* test). (H-J) Normalized translation rates as measured by incorporation of methionine analog HPG over time and evaluated by flow cytometry. Each data point represents the slope of HPG incorporation for at least 3 time points using median fluorescence intensity from an independent experiment. LM2 cells from (E) following 3 days of doxycycline (1 μg/mL) induction (H), LM2 cells following 3 days of control or OICR-9429 treatment at 20 μM (I), and LM2 cells following 3 days of MS67N or MS67 treatment at 2.5 μM (J) were tested. (n=3, one sample t-test). *p<0.05; **p<0.01; ***p<0.001; ****p<0.0001.

We then performed Enrichr analysis on both the up- or down-regulated DEGs and found that the most enriched and significant gene ontology in the shared down-regulated DEGs was cytoplasmic ribosomal proteins (**Figure 3C, Figure 3-figure supplement 1A, Table S2**). The combined score for this enrichment was 18-fold higher than the next enriched ontology for the shared down-regulated DEGs, demonstrating the significance of this WDR5 regulated pathway (**Figure 3C, Table S2**). Notably, among the 474 down-regulated DEGs in LM2 cells, 51 (11%) encoded for ribosomal protein (RP) genes (**Figure 3D**). A similar enrichment pattern was observed in BrM3 and BoM cells (**Figure 3-figure supplement 1B, 1C**). After inducing WDR5 knockdown with both hairpins for 3 days in LM2 cells, we confirmed down-regulation of all the tested RP genes. These included the top two down-regulated RPs, *RPL7* and *RPL31*, which were consistently reduced by a ∼50% (**Figure 3E**). We next tested whether WDR5 targeting with either OICR-9429 or MS67 would have similar effects on gene expression. Treatment of LM2 and MDA-MB-453 cells with 10 or 20 μM of OICR-9429 for 3 days decreased *RPL7* and other RP genes as predicted (**Figure 3F, Figure 3-figure supplement 1D**). We next evaluated the gene-regulatory effect of MS67 which is a more effective inhibitor of WDR5. MS67 treatment at 2.5 μM significantly downregulate several RP genes whereas OICR-9429 and MS67N did not have an effect at these lower concentrations (**Figure 3G, Figure 3-figure supplement 1E**). As down-regulation of RP genes expression implies a decrease in ribosome biogenesis, we also measured protein translation rates in TNBC cells where WDR5 was pharmacologically or genetically blocked. Accordingly, WDR5 silencing impaired global protein translation rates (**Figure 3H, Figure 3-figure supplement 1F**). OICR-9429 treatment or MS67-mediated degradation also caused decreases in protein translation in both LM2 and MDA-MB-453 cell lines (**Figure 3I, 3J, Figure 3-figure supplement 1G**). Taken together, our data demonstrates that either genetic or pharmacological inhibition of WDR5 can suppress RP gene expression and global translation in breast cancer cells.

### The WBM binding sites are required for WDR5-dependent cell growth and ribosomal protein gene expression

We next sought to identify which of WDR5’s multiple molecular functions is required for RP gene expression and breast cancer cell growth. WDR5 is canonically part of the mammalian KMT2A complex, which also consists of WRAD proteins (**W**DR5, **R**BBP5, **A**SH2L, and **D**PY30). Only KMT2A and RBBP5 interact directly with WDR5 and the complex has recently been elucidated by cryo-electron microscopy (Park et al., 2019) (**Figure 4A**). WDR5 is a donut-shaped protein with two important binding pockets, WIN and WBM (Guarnaccia & Tansey, 2018) (**Figure 4A and 4B**). KMT2A binding to the WIN site can be disrupted by point mutation F133A on WDR5 (Guarnaccia et al., 2021; Patel et al., 2008). On the other hand, RBBP5 and c-MYC have been shown to bind at the WBM site, which can be disrupted by point mutations N225A and V268E (Guarnaccia et al., 2021; Thomas, Wang, et al., 2015). In addition to the F133A, N225A, and V268E mutants, WDR5 mutants K7Q and 1-25Δ, were recently shown to specifically impact ciliogenesis (Kulkarni et al., 2018).

**Figure 4.**
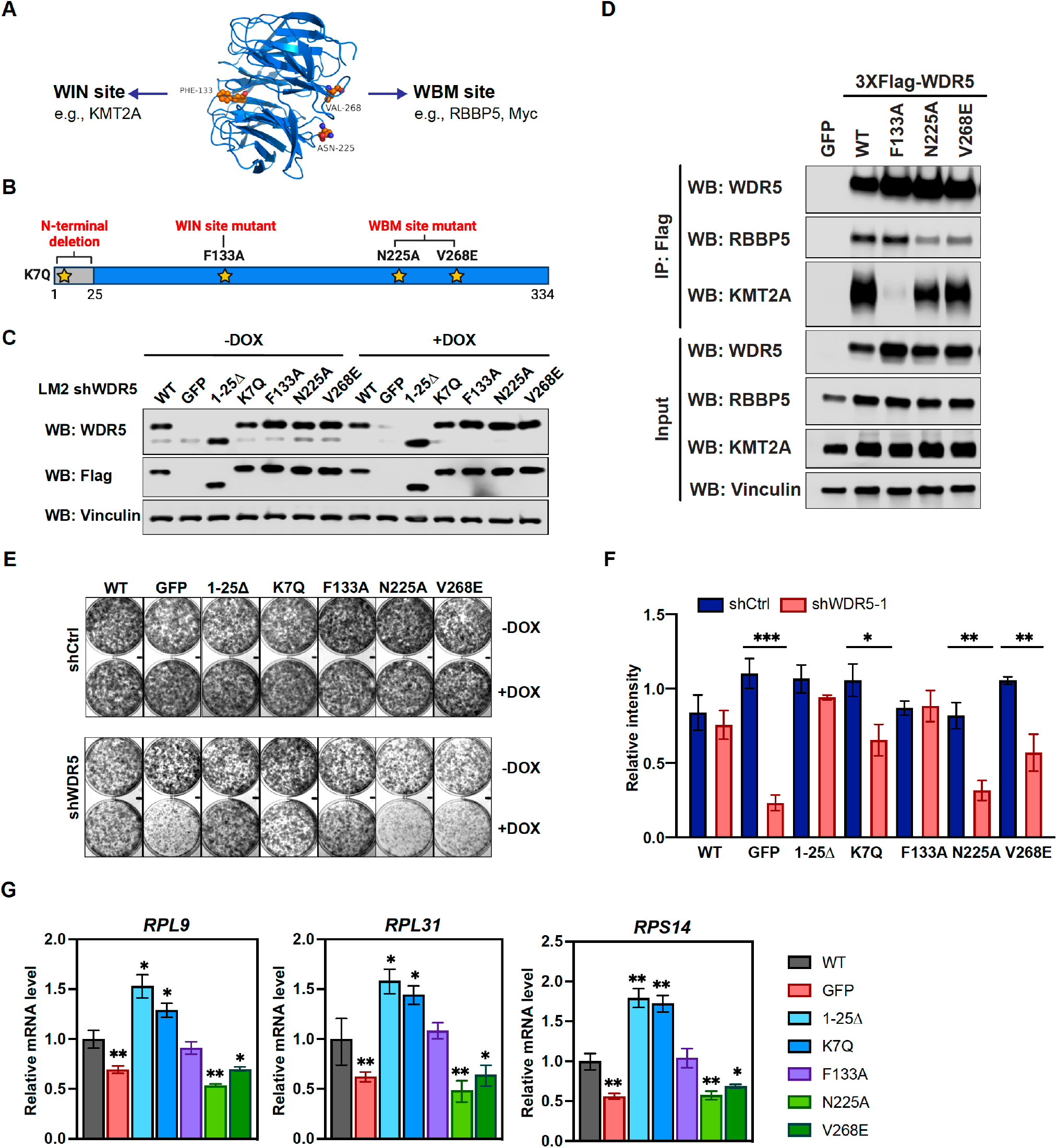
The WBM binding sites are required for WDR5-dependent cell growth and ribosomal protein gene expression. (A) WDR5 protein structure and key residues in the WIN and WBM sites that interact with binding partners. (B) Schematic of WDR5 with indicated mutation sites. (C) Western blot analysis of indicated proteins in LM2 inducible shWDR5 cells over-expressing WT WDR5 or WDR5 mutants. Cells were collected after 3 days of control or doxycycline (1 μg/mL) induction. (D) Western blot analysis of the indicated proteins after immunoprecipitation using anti-Flag antibody in the LM2 shWDR5 cells over-expressing GFP, WT WDR5, or WDR5 mutants. (E-F) Colony formation assays of cells expressing GFP, WT WDR5, or WDR5 mutants in inducible shControl (shCtrl) or shWDR5 cell lines after 9 days of control or doxycycline (1 μg/mL) treatment. Representative images (E) and quantification (F) are shown. Doxycycline treated wells were compared to their respective controls for each cell line (n=3, unpaired two-side Student’s *t* test). (G) RT-qPCR analysis of the indicated mRNAs in LM2 from (E) induced with doxycycline (1 μg/mL) for 3 days. Significance determined by comparing each treatment to WT control (n=4, unpaired two-side Student’s *t* test). *p<0.05; **p<0.01; ***p<0.001; ****p<0.0001. For gel source data, see Figure 4- source data1-2.

Based on this information, we performed a structure function analysis of WDR5 by constitutively expressing shRNA-resistant wild-type (WT) WDR5 or the aforementioned WDR5 mutants with C-terminal 3XFlag-tag in LM2 cells, where endogenous *WDR5* was concomitantly silenced. Following DOX induction, we confirmed ectopic WDR5 (mutant or wild-type) expression in the indicated mutant cell lines, whereas endogenous WDR5 levels were significantly repressed (**Figure 4C**). Using co-immunoprecipitation assays, we observed that the F133A but not N25A or V268E mutations abrogate the binding of WDR5 to KMT2A. Alternatively, mutants N225A and V268E but not F133A reduced the binding of WDR5 to RBBP5 by more than 50% as expected (**Figure 4D**). We next determined which WDR5 interacting site is required for cell growth. Consistently, shWDR5 cells expressing GFP control had severely impacted colony formation while expression of WT WDR5 rescued this growth defect (**Figure 4E and 4F**). The N-terminal mutant 1-25Δ or K7Q have either a similar or slightly lower ability to rescue WDR5 dependent cell growth, respectively. Surprisingly, the WIN site mutant F133A was able to rescue the colony formation phenotype, while neither N225A nor V268E effectively rescued cell growth. These results suggest that the WBM but not the WIN binding ability of WDR5 is required for WDR5-dependent growth of TNBC cells.

We next tested whether the different WDR5 mutants affected WDR5 binding to the promoter of RP genes and alter H3K4me3 levels in LM2 cells. ChIP-qPCR analysis showed that the F133A mutant binds to the promoter of *RPL7* and *RPL31* less efficiently, while N225A and V268E mutants bind chromatin similarly as the WT protein (**Figure 4-figure supplement 1A**). Surprisingly, all mutants maintained a similar level of H3K4me3 (**Figure 4-figure supplement 1B**), suggesting that WDR5 binding to chromatin is not required for maintaining H3K4me3 at the RP gene promoters tested in this context. More importantly, the N-terminal and F133A mutants rescued the expression of *RP* genes, whereas *the* N225A and V268E mutants did not **(Figure 4G)**. Altogether, these data suggest that, in LM2 cells, WBM but not WIN binding by WDR5 is important for the maintenance of RP gene expression.

### Metastatic cell growth and lung colonization do not require the KMT2 complex components

The surprising observation that WIN binding by WDR5 is dispensable for breast cancer cell growth prompted us to directly test the requirement for the canonical KMT2 complex components in LM2 cells (**Figure 5A**). We first confirmed efficient knockdown of 7 complex components (KMT2A, RBBP5, DPY30, HCFC1, CXXC1, WDR82, BOD1L1), each with two independent shRNA after 3 days of DOX induction (**Figure 5B and 5C**). We first directly asked if KMT2A is required for LM2 cell growth and lung metastasis as KMT2A is a catalytic subunit of the H3K4 methyltransferase complex and was seemingly depleted in our screen (**Figure 1B, 1C**). Silencing KMT2A with multiple shRNAs did not reproducibly affect *in vitro* colony formation and *in vivo* lung metastasis growth (**Figure 5D-F**). This result is also consistent with the phenotype observed from the F133A mutant and suggests that KMT2A is dispensable for WDR5-dependent cell growth. We next assessed whether RBBP5, DPY30, and HCFC1 are required for RP expression and cell growth. RP gene expression was not changed after knockdown of RBBP5, DPY30, or HCFC1 (**Figure 5G**). Furthermore, most of these KMT2 complex components are not required for the growth of LM2 cells (**Figure 5H and 5I**). The exception was upon knockdown of HCFC1 which resulted in a 40% decrease in colony formation (**Figure 5H and 5I**), likely due to the role of HCFC1 in cell cycle control (Antonova et al., 2019; Xiang et al., 2020) While RBBP5, DPY30 and HCFC1 are common to the KMT2 complexes, CXXC1 and WDR82 are distinct to the SET1A/B complexes and BOD1L1 is specific to the SET1B complex (**Figure 5A**). Thus, we asked whether WDR5 regulates the growth phenotype specifically through SET1A/B complexes by perturbing CXXC1, WDR82, or BOD1L1. However, knockdown of these components also did not decrease RP gene expression and colony formation (**Figure 5G-I**). Therefore, the KMT2 complexes are not the major effectors of WDR5 dependent metastatic cell growth in the LM2 model.

**Figure 5.**
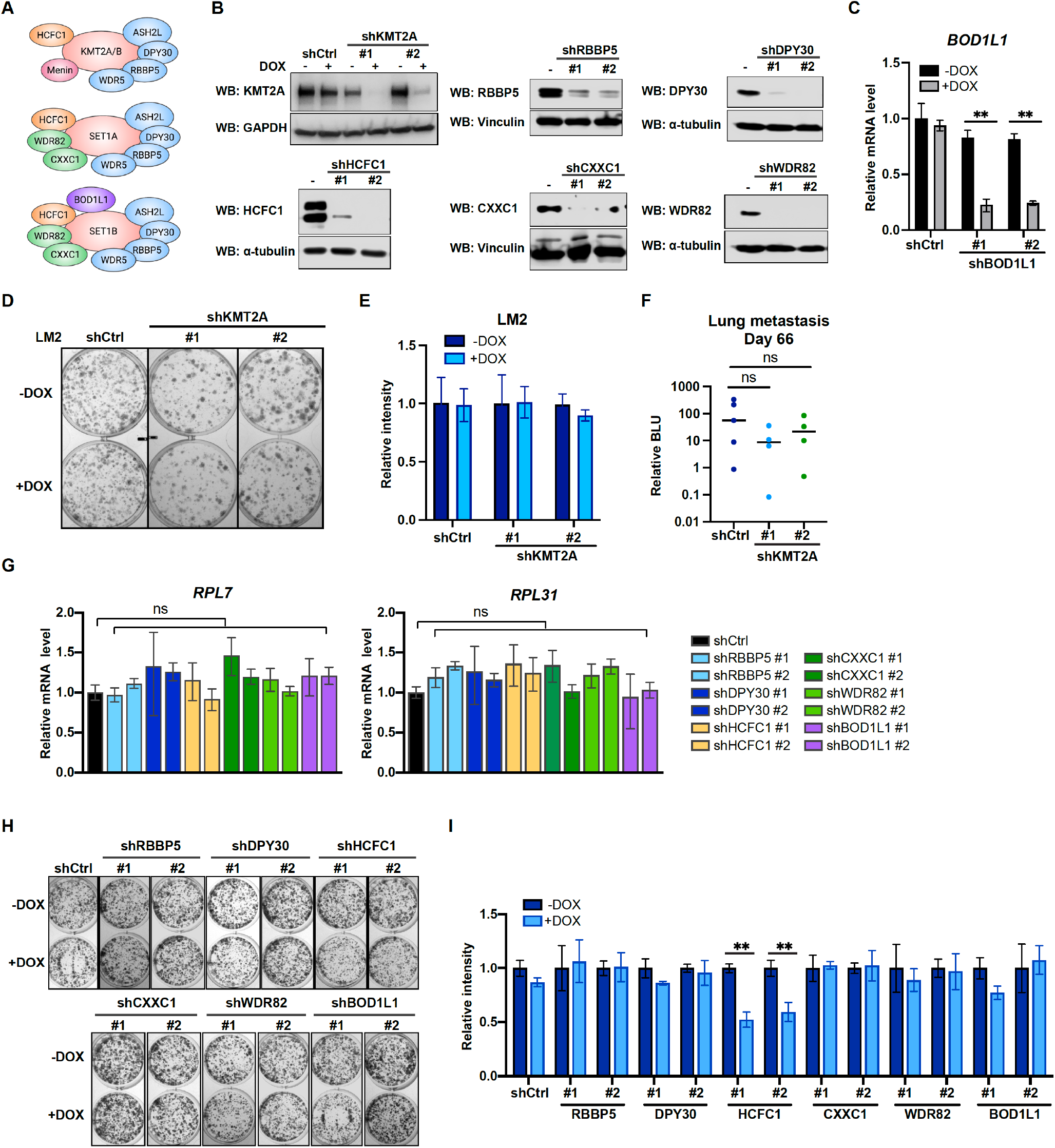
Metastatic cell growth and lung colonization do not require KMT2 complex components. (A) Schematic of subunit composition of several KMT2 complexes. (B) Western blot analyses of the indicated proteins in LM2 cells transduced with inducible shRNA targeting KMT2A, RBBP5, DPY30, HCFC1, CXXC1, and WDR82. Cells were collected after 3 days of doxycycline (1 μg/mL) treatment. (C) RT-qPCR analysis of *BOD1L1* in LM2 cells transduced with two independent hairpins targeting *BOD1L1*. Cells were collected after 3 days of doxycycline treatment (n=4, unpaired two-side Student’s *t* test). (D-E) Colony formation assay of LM2 shCtrl or shKMT2A cells (shKMT2A-1and shKMT2A-2) after 9 days of treatment with control or 1 μg/mL doxycycline. Representative images (D) and quantification (E) are shown (n=3, unpaired two-side Student’s *t* test). (F) Normalized bioluminescence signals of lung metastasis at day 66 of mice injected intravenously with LM2 cells from (B) and kept under doxycycline chow. The data represent mean ± SEM. Significance determined using unpaired two-tailed Mann-Whitney test. (G) RT-qPCR analysis of LM2 cells transduced with the indicated inducible shRNAs. Cells were collected after 3 days of doxycycline treatment. Significance determined by comparing each treatment to shCtrl. (H-I) Colony formation assay of LM2 cells from (G) after 9 days of either control or doxycycline (1 μg/mL) treatment. Representative images (H) and quantification (I) are shown (n=3, unpaired two-side Student’s *t* test). *p<0.05; **p<0.01; ***p<0.001; ****p<0.0001. For gel source data, see Figure 5- source data1.

### Inhibition of WDR5 and mTOR cooperatively reduces translation and TNBC growth

Hyperactivation of growth signaling pathways can increase protein synthesis and inhibition of translation is being actively explored as a therapeutic avenue for cancer (Bhat et al., 2015; Grzmil & Hemmings, 2012). Several mTOR inhibitors have been approved or are being tested in clinal trials, including the first generation mTOR inhibitors, everolimus and temsirolimus, and the second generation mTOR inhibitor, OSI-027 (Zheng & Jiang, 2015). Everolimus and temsirolimus are rapalogs that allosterically inhibit mTORC1, while OSI-027 is an ATP-competitive inhibitor that inhibits both mTORC1 and mTOCR2 (Zheng & Jiang, 2015). Everolimus has been approved to treat postmenopausal women with advanced hormone receptor positive, HER2 negative breast cancer in combination with an aromatase inhibitor exemestane (Baselga et al., 2012). Because cancer cells could develop resistance to inhibitors of protein translation and this class of drugs may not directly cause cell death (Rozengurt et al., 2014; Zheng & Jiang, 2015), identifying other regimens which synergize with mTOR inhibitors is warranted.

Interestingly, during the course of titrating mTOR inhibitors in our cell model, we noted that the treatment with OSI-027 or everolimus alone caused an up-regulation of RP gene expression (**Figure 6-figure supplement 1A-D**), which may be due to an adaptive feedback effect on proteostasis following mTOR inhibition. Importantly, treatment with the WDR5 inhibitor OICR-9429 partially or completely blocked this adaptive induction of RP genes (**Figure 6-figure supplement 1A, 1B**). Moreover, while mTOR inhibitors were confirmed to down-regulate phosphorylated S6 protein kinase (S6K) and translation initiation factor 4E binding protein 1 (4E-BP1), WDR5 inhibition reduced RP genes expression and translation independently of this signaling pathway (**Fig 6A, Figure 6-figure supplement 1E, 1F**).

**Figure 6.**
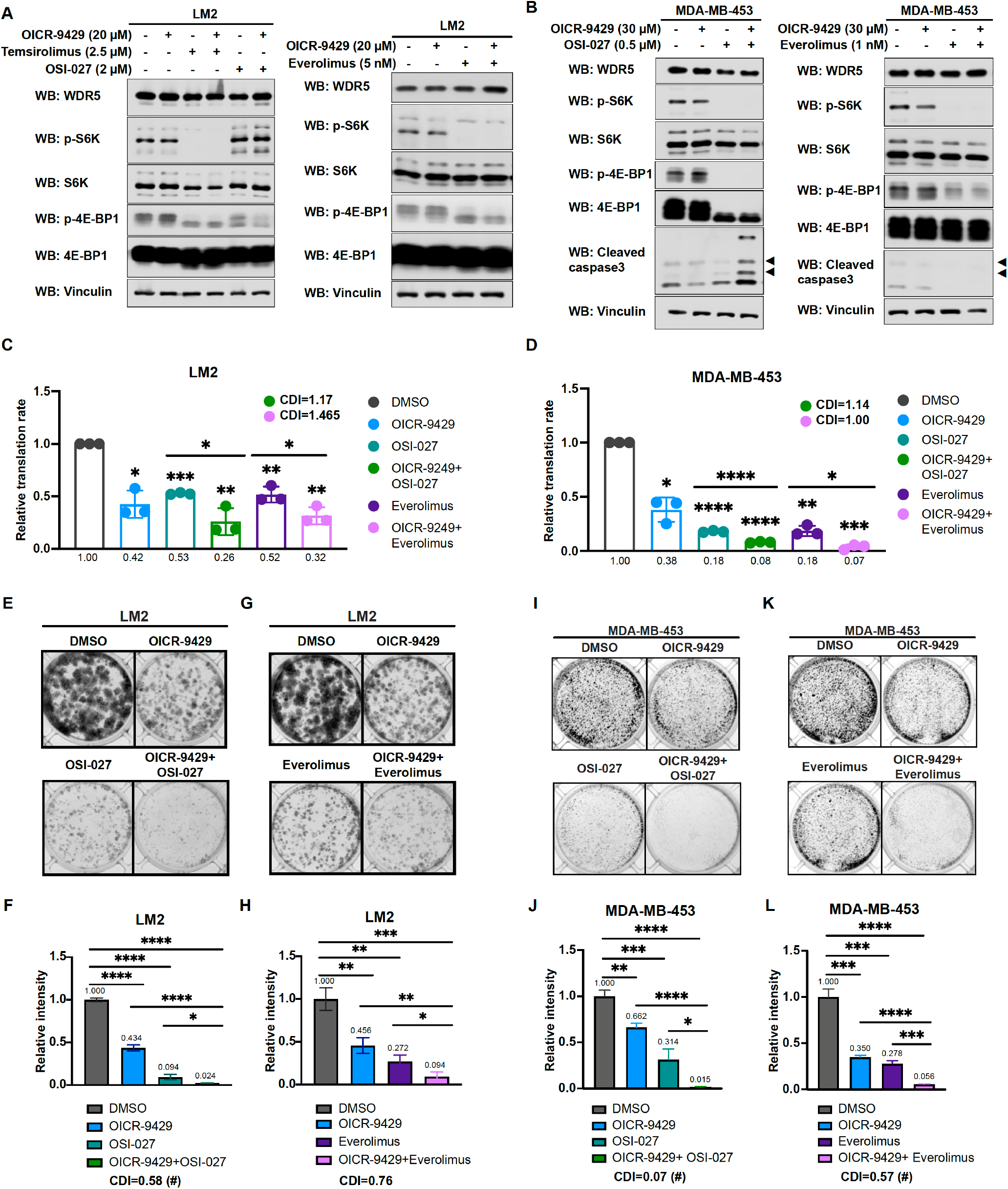
Inhibition of WDR5 and mTOR cooperatively reduces translation and TNBC growth. (A) Western blot analysis of the indicated proteins in LM2 cells with or without 20 μM OICR-9429 in combination with 3 days of control, 2 μM OSI-027, 2.5 μM temsirolimus, or 5 nM everolimus treatment. (B) Western blot analysis of the indicated proteins in MDA-MB-453 with or without 30 μM OICR-9429 in combination with 3 days of control, 0.5 μM OSI-027, or 1 nM everolimus treatment. (C) Normalized translational rates of LM2 cells from (A) (n=3, one sample t-test). (D) Normalized translational rates of MDA-MB-453 cells from (B) (n=3, one sample t-test). (E-F) Colony formation assay of LM2 with or without 20 μM OICR-9429 in combination with control or 2 μM OSI-027 treatment for 8 days. Representative images (E) and quantification (F) are shown. (G-H) Colony formation assay of LM2 cells with or without 20 μM OICR-9429 in combination with control or 5 nM everolimus treatment for 8 days. Representative images (G) and quantification (H) are shown. (I-J) Colony formation assay of MDA-MB-453 with or without 30 μM OICR-9429 in combination with control or 0.5 μM OSI-027 treatment for 10 days. Representative images (I) and quantification (J) are shown. (K-L) Colony formation assay of MDA-MB-453 with or without 30 μM OICR-9429 in combination with control or 1 nM everolimus treatment for 10 days. Representative images (K) and quantification (L) are shown (n=3, unpaired two-side Student’s *t* test). *p<0.05; **p<0.01; ***p<0.001; ****p<0.0001. Calculation of coefficients of drug interaction (CDI) is described in materials and methods section. Significant synergy is labeled with (#). For gel source data, see Figure 6- source data1-2.

Based on these results, we postulated that the inhibition of WDR5 and mTOR could cooperatively decrease TNBC protein translation, cell growth, and survival. As such we first treated LM2 and MDA-MB-453 cells for 3 days using OICR-4129 or the three mTOR inhibitors, everolimus, temsirolimus, and OSI-027. The levels of phosphorylated S6 protein kinase (S6K) and translation initiation factor 4E binding protein 1 (4E-BP1) were decreased in both cell lines after everolimus or temsirolimus treatment, while OSI-027 treatment only showed strong inhibition of mTOR signaling in the MDA-MB-453 cells (**Figure 6A and 6B**). Similar mTOR signaling inhibition was observed in LM2 cells expressing shWDR5, confirming that mTOR regulation is independent of WDR5 (**Figure 6-figure supplement 1F**). Next, we compared the global translation rates of LM2 or MDA-MB-453 cells, when WDR5 was inhibited genetically or pharmacologically in combination with mTOR inhibitors OSI-027 or everolimus. Overall translation was decreased when combining WDR5 inhibition or WDR5 silencing with mTOR inhibition (**Figure 6C, 6D, Figure 6-figure supplement 1G**). This combinatorial effect on protein translation correlated with an additive inhibition of clonogenic outgrowth (**Figure 6-figure supplement 1H-1K**). Moreover, both OSI-027 and everolimus act synergistically with WDR5 inhibition in both LM2 and MDA-MB-453 cells (**Figure 6E-6L**), while temsirolimus showed additive effect in LM2 cells (**Figure 6-figure supplement 1L, 1M**). We next tested the effects of MS67-mediated WDR5 degradation in combination with mTOR inhibition. MS67 treatment alone also did not affect mTOR signaling (**Figure 7A and 7B**). Interestingly, MS67 acts synergistically with both OSI-027 and everolimus in inhibiting translation in MDA-MB-453 cells (**Figure 7C**). Furthermore, we found that 5 µM MS67 is more effective than 20 µM OICR-9429 at inhibiting colony outgrowth when combined with either OSI-027 or everolimus in LM2 cells (compare **Figure 6E-H** with **Figure 7D-G**). Importantly, we found that OSI-027 had better synergistic effects with WDR5 inhibition when compared to everolimus, suggesting that mTORC2 could be critical for clonogenic outgrowth in the context of WDR5 inhibition. Moreover, we observed increased cleaved-caspase 3 level in the combined treatment group in MDA-MB-453 cells (**Figure 6B**), suggesting the combination of WDR5 inhibition and OSI-027 induces apoptosis.

**Figure 7.**
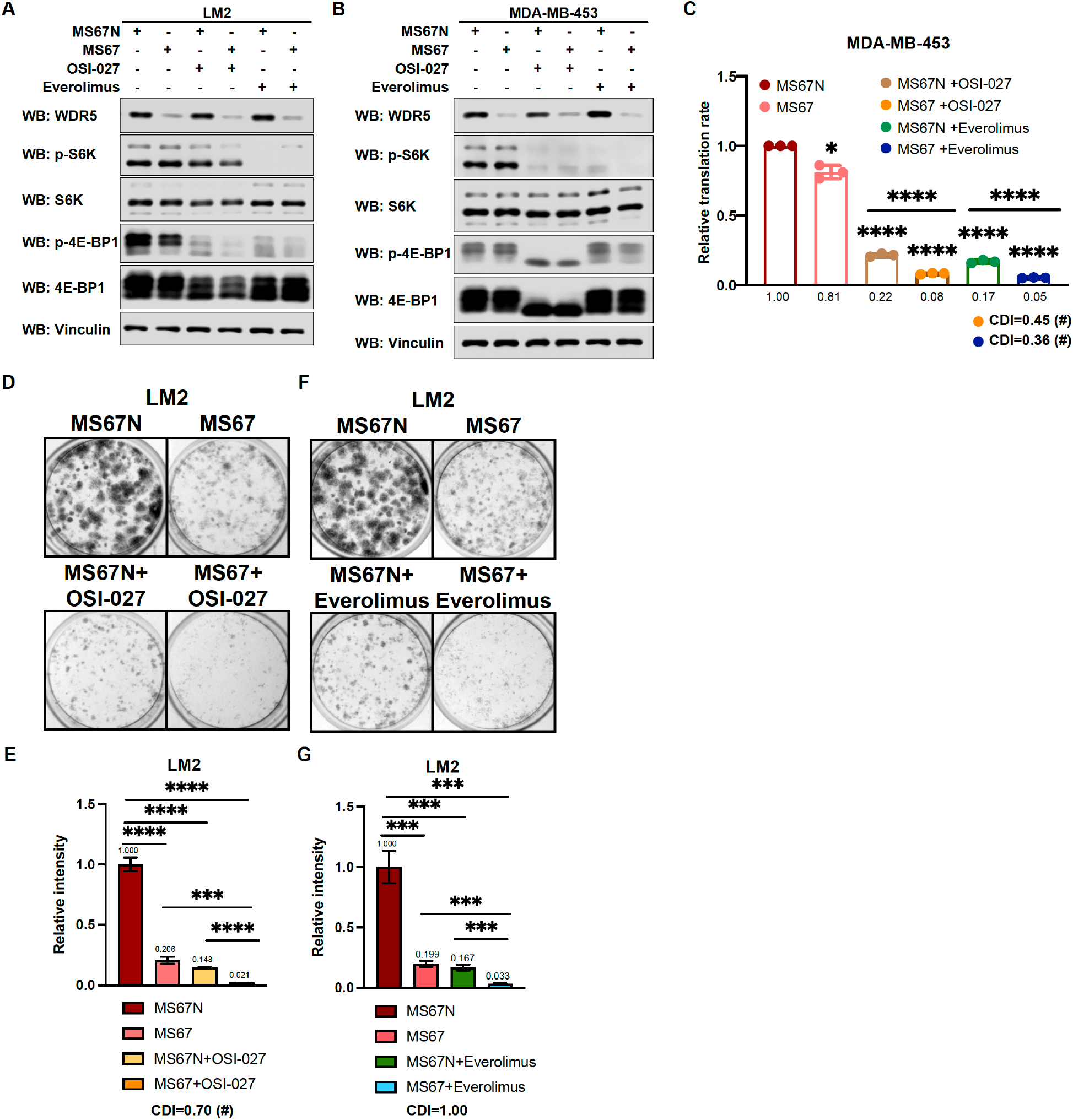
MS67-mediated WDR5 degradation and mTOR inhibition cooperatively reduces translation and TNBC growth. (A) Western blot analysis of the indicated proteins in LM2 with 2.5 μM MS67N or MS67 in combination with 3 days of control, 2 μM OSI-027, or 5 nM everolimus treatment. (B) Western blot analysis of the indicated proteins in MDA-MB-453 with 2.5 μM MS67N or MS67 in combination with 3 days of control, 0.5 μM OSI-027, or 1nM everolimus treatment. (C) Normalized translational rates of MDA-MB-453 cells from (B) (n=3, one sample t-test). (D-E) Colony formation assay of LM2 cells with 5 μM MS67N or MS67 in combination with control or 2 μM OSI-027 for 9 days. Representative images (D) and quantification (E) are shown. (F-G) Colony formation assay of LM2 cells with 5 μM MS67N or MS67 in combination with control or 5 nM everolimus for 9 days. Representative images (F) and quantification (G) are shown (n=3, unpaired two-side Student’s *t* test). *p<0.05; **p<0.01; ***p<0.001; ****p<0.0001. Calculation of coefficients of drug interaction (CDI) is described in materials and methods section. Significant synergy is labeled with (#). For gel source data, see Figure 7- source data1-2.

Collectively, our data identified WDR5 mediated protein translation as a potential vulnerability, which could be therapeutically leveraged in TNBC cells treated with first-generation or second-generation mTOR inhibitors.

## Discussion

Epigenetic aberrations contribute to multiple steps of tumor initiation, cancer invasion and migration, and tumor outgrowth through a wide spectrum of mechanisms (Blair & Yan, 2012; Chen & Yan, 2021). Moreover, recent efforts have led to the development of multiple pharmacological agents designed to target epigenetic and chromatin modifying proteins in cancer (Ahuja et al., 2016; Lu et al., 2020; Zhou et al., 2020). Nevertheless, it is unclear how such agents can be leveraged therapeutically as single agents or in combination, particularly for the treatment of breast cancers. In this study, we performed an *in vivo* functional screen of epigenetic factors to identify WDR5 as being required for metastatic breast cancer growth. Intriguingly, WDR5 regulates ribosomal gene expression independent of its H3K4 methylation activity but through its WBM domain to mediate translation rate and cell growth. WDR5 inhibition or degradation suppresses translation and growth of breast cancer cells, alone or in combination with mTOR inhibitors. These results indicate that WDR5 promotes breast cancer growth and metastasis through regulating translation.

WDR5 is best known for its role in the KMT2 complexes, which promote transcription through H3K4 methylation (Wysocka et al., 2005). Unexpectedly, our structure function studies using the F133A WDR5 mutant, suggest that KMT2 binding may not be critical for WDR5 mediated ribosomal gene expression and cell growth by metastatic TNBC cells. Consistently, depletion of several other components of the KMT2 complex did not affect the fitness of metastatic TNBC cells. These results suggest that WDR5 regulates translation and growth through KMT2 enzymatic activity-independent function.

WDR5 can also be recruited to the NSL complex with the acetyltransferase MOF, and WDR5 directly interacts with the subunit KANSL1 and KANSL2 through WIN and WBM sites, respectively (Dias et al., 2014). The interaction of KANSL1 to WDR5 is important for efficient targeting of NSL complex to the promoter of target genes (Dias et al., 2014). Therefore, it is likely that the NSL complex does not contribute to these phenotypes as WIN site WDR5 mutant F133A did not show a defective growth phenotype in this context. Alternatively, WDR5 likely regulates the phenotype described herein through a non-canonical function, such as its known ability to recruit the transcription factor MYC (Thomas, Wang, et al., 2015). WDR5 were previously shown to directly interact with MYC through WBM site and facilitate the recruitment of MYC to chromatin (Thomas, Wang, et al., 2015). This is consistent with our findings that WBM site mutants of WDR5 are unable to rescue the growth defect caused by WDR5 loss. Notably, the association of MYC to its target genes is disrupted when the WBM site is mutated (Thomas, Wang, et al., 2015).

A recently published study implicates WDR5 in maintaining metastatic outgrowth via trimethylation of H3K4 on the promoters of specific target genes including *TGFB1*, which enhances epithelial to mesenchymal transition (EMT) (Punzi et al., 2019). Alternatively, by using a genome-wide approach and multiple TNBC cell line models, we did not observe alterations in EMT, which may be context specific (Figure 3C and S3A). Conversely, we demonstrated a conserved and broad role for WDR5 in controlling ribosomal protein (RP) gene expression (including *RPL32*, *RPL34*, *RPS14*, and *RPS6*) in a manner that is independent of KMT2 and H3K4me3 at the promoters of RP genes. Therefore, the primary role of WDR5 may be to regulate proteostasis in TNBC cells. As aberrant protein translation affects multiple features of malignant cells, targeting WDR5 would be effective in treating both early or late stages of breast cancer (Grzmil & Hemmings, 2012). Consistent with this idea, knockdown of WDR5 independently decreases primary tumor growth and lung metastasis *in vivo*. Future studies will be needed to elucidate how WDR5-dependent protein translation contributes to the different steps of breast cancer progression, dissemination, and colonization.

The regulation of proteostasis and targeting protein translation in particular, are potential therapeutic vulnerabilities of cancer cells. Interestingly, we demonstrated that the regulation of RP gene expression and protein translation could be inhibited by using the WDR5 inhibitor OICR-9429 or WDR5 degrader MS67, consistent with our genetic approach with WDR5 gene knockdown. Proteolysis targeting chimeras (PROTACs) are hetero-bifunctional small molecules that can recruit desired target protein to the E3 ubiquitin ligase complex for proteasomal degradation (Paiva & Crews, 2019). Multiple PROTAC degraders have entered clinical trials for cancer treatment (He et al., 2020). Here, we leveraged the newly designed WDR5 degrader to test its efficacy in WDR5 degradation in breast cancer cells. In fact, the WDR5 degrader MS67 showed superior effect than the WDR5 inhibitor OICR-9429. MS67 led to WDR5 degradation within 4 hours and is reversible after withdrawal of drug treatment (Yu et al., 2021), allowing for temporal control of WDR5 targeting. Unlike small molecule inhibitors, PROTAC molecules can be reused within the cells, which would lower the required concentration for drug treatment. Additionally, PROTAC is able to degrade the entire protein in the cells, which could overcome some potential drug resistant mechanisms. Our results thus suggest that WDR5 degradation is a potential therapeutic strategy to inhibit metastatic progression in breast cancer.

Finally, we discovered that multiple mTOR inhibitors can act synergistically with WDR5 targeting. In addition, we show that the second generation mTOR inhibitor, OSI-027, which targets both mTORC1 and mTORC2, works better than first generation inhibitor, everolimus, when treated in combination with WDR5 targeting. Both mTOR inhibition or WDR5 degradation can inhibit translation but through different mechanisms. mTOR integrates survival signals with protein synthesis. As translation initiation is initially repressed upon mTOR inhibition, negative feedback loops can cause aberrant stimulation of upstream signaling via AKT activation, which may diminish the effect of mTOR inhibitors (Rozengurt et al., 2014; Zou et al., 2020). We also observed up-regulation of RP gene expression after mTOR inhibitor treatment, suggesting that epigenetic activation of ribosomal genes may be another compensatory response to mTOR inhibition. Importantly, we demonstrated that WDR5 inhibition is able to counteract this feedback activation of RP genes. Altogether, our study provides molecular and cell biological evidence that WDR5 is an important epigenetic mediator of protein translation and that this distinct function of WDR5 may be leveraged for treatment of TNBC.

## Materials and methods

### Antibodies and chemicals

For co-IP and western blots, the following antibodies were obtained commercially: mouse anti-Flag (M2, F1804), mouse anti-vinculin (V9131), and mouse anti-tubulin (T5168) (Sigma, St. Louis, MO); rabbit anti-WDR5 (#13105), rabbit anti-RBBP5 (#13171), rabbit anti-KMT2A/MLL1-C (#14197), rabbit anti-CXXC1(#12585), rabbit anti-HCFC1 (#69690), rabbit anti-WDR82 (#99715), rabbit anti-phospho p70 S6K (Thr389) (#9205), rabbit anti-p70 S6K (#9202), rabbit anti-phospho 4E-BP1 (Thr37/46) (#2855), rabbit anti-4E-BP1 (#9644), and anti-Cleaved caspase 3 (#9661) (Cell Signaling Technology, Danvers, MA); rabbit anti-DPY30 (A304-96A) (Bethyl Laboratories, Montomery, TX).

For drug treatment experiments, WDR5 inhibitor OICR-9429 (Sigma, SML1209 and Cayman Chemical, #16095), and mTOR inhibitors, OSI-027 (Cayman Chemical, #17379), everolimus (Cayman Chemical, #11597), and temsirolimus (Cayman Chemical, #11590) were used. Compounds of WDR5 degrader MS67 and negative control MS67N were synthesized in Jian Jin’s lab.

### Plasmids and virus generation

Frozen bacterial stocks harboring the shRNA library were generated by the Westbrook lab. pGIPZ plasmid harboring hairpins and barcodes were digested with Xho I and Mlu I and sub-cloned into the pINDUCER10 plasmid. The list of hairpin sequences is available in **Table S3**. For cloning of the WDR5 mutants, BP cloning primers were designed against p3XFlag-CMV-14-WDR5. Two-step PCR was performed to generate shRNA resistant mutant WDR5. Briefly, two sets of primers were designed such that they overlap at the site of mutagenesis. The product from the PCR was then used for BP (Thermo Fisher, # 11789020) or LR (Thermo Fisher, #11791020) reaction into pDONR-211 or pLenti-PSK-hygro-DEST. p3XFlag-CMV-14-WDR5 was a gift from Debu Chakravarti (Addgene #59974). A list of cloning oligos is available in **Table S4**.

For virus generation, HEK293T cells were transfected with 1.2 µg each of VSV-G, TAT, RAII, and HyPM packaging plasmids along with 11.2 µg of lentiviral plasmid. OptiMEM and TransIT-293 Transfection Reagent (Mirus, MIR2700) were used following manufacturer protocol. Viruses were collected at 48h and 72h, filtered through a 0.45 μm filter.

### Cell culture and stable cell lines generation

MDA-MB-231 and its metastatic derivatives, MDA-MB-231-LM2 (LM2), MDA-MB-231-BoM (BoM) and MDA-MB-231-BrM3 (BrM3) breast cancer cells and HEK293T cells were cultured in Dulbecco’s Modified Eagle Medium supplemented with 10% fetal bovine serum and 100 U/mL penicillin, and 100 µg/mL streptomycin. HCC1143, MDA-MB-453, MCF7, T47D, MDA-MB-361, UACC893, BT474, SKBR3, and 4T1 breast cancer cells were cultured in RPMI1640 supplemented with 10% fetal bovine serum, 100 U/mL penicillin, and 100 µg/mL streptomycin. Cells were periodically tested for mycoplasma contamination and authenticated using short tandem repeat profiling.

For generation of cell lines, viruses harboring pINDUCER10-puromycin or pINDUCER10-blasticidin constructs were titrated using the target cell lines. Cells were infected at an MOI of 1 and selected using either 0.8 µg/mL puromycin or 10 µg/mL blasticidin. For generation of cell lines harboring WDR5 mutants, optimal viral dose was determined empirically by western blot visualization to assess equal expression of WDR5 across mutant cell lines. LM2 cells with re-introduction of WDR5 mutants were selected with 800 µg/mL hygromycin.

### Minipool generation for *in vitro* and *in vivo* screening

Minipools were created by equally mixing 8-10 individual LM2 cell lines harboring pINDUCER10 hairpins targeting each epigenetic modifier together with two LM2 positive control cell lines (shBUD31 and shSAE2) and two negative control cell lines (shCHEK1 and shSTAMBP). For *in vitro* screening, minipool cells were plated into 10-cm dishes with or without 1 μg/mL of doxycycline. A portion of minipool cells were collected as day 0 samples as the controls. Every two days the cells were pelleted all samples were proceeded to gDNA isolation and gDNA qPCR. For *in vivo* screening, 5×10^5^ minipool cells were injected into nude mice through tail vein. Lung metastases were monitored weekly with *in vivo* live imaging. At the end point, the mice were sacrificed and the lung tissue was harvested for gDNA isolation and gDNA qPCR. For the screening readout analyses, all qPCR results were normalized to the value from day 0. The fold change was obtained from +DOX/-DOX for both *in vitro* and *in vivo* screen.

### Animal studies

Female Athymic Nude-*Foxn1^nu^* immunodeficient (6-8 weeks old) mice (Envigo) were used for lung-metastasis experiments with human cell lines. For *in vivo* screening 5×10^5^ cells were injected via tail vein in 0.1 ml saline. For WDR5 *in vivo* validation experiment, cells were treated with doxycycline for 3 days prior to injection and 2×10^5^ cells were injected via tail vein in 0.1 ml saline. Mice were placed on doxycycline chow (Envigo, TD.01306) 5 days prior to injection. All the *in vivo* metastasis signals, including lung metastasis and whole-body metastasis, were monitored by weekly bioluminescence imaging with an IVIS system coupled to Living Image acquisition and analysis software (Xenogen). Luminescence signals were quantified at the indicated time points as previously described. Values of luminescence photon flux of each time point were normalized to the value obtained immediately after xenografting (day 0).

For mammary fat pad tumor assays, control and shWDR5-1 LM2 cells (1×10^6^) were resuspended in 0.1 mL of saline and matrigel (corning #356231) mix, and then injected into mammary fat pad (the 4^th^ mammary glands) of NOD-SCID mice (6 weeks old). Tumor were monitored every 7 days by measuring the tumor length (L) and width (W). Tumor volume was calculated as V=L×W^2^/2. Mice were euthanized when primary tumors reached 1,000 mm^3^. All animal procedures were approved by the Institutional Animal Care and Use Committee of Yale University.

### Lung tissue harvest and gDNA isolation

Mice were sacrificed and whole body perfused with 10 mL of PBS. For gDNA isolation, the harvested lungs were placed into a microcentrifuge tube and snap-frozen with liquid nitrogen. The frozen tissues were then placed into an aluminum block on dry ice. Each tube of the lung tissue was allowed to thaw enough for further mincing with surgical scissors, and then refrozen by dipping them in liquid nitrogen bath. This process was repeated 2-3 times until no visible tissue chunk was observed. 60 mg of homogenized tissue was then aliquoted out and processed with the QIAmp DNA mini kit (Qiagen 51304) following manufacturer’s protocols.

### Western blot and Co-immunoprecipitation (Co-IP)

Cells were lysed in 1X high salt lysis buffer (50 mM Tris-HCl pH 8, 320 mM NaCl, 0.1 mM EDTA, 0.5% NP-40, 10% glycerol) or RIPA buffer (50 mM Tris-HCl pH 7.4, 150 mM NaCl, 1 mM EDTA, 1% Triton X-100, 1% sodium deoxycholate, 0.1% SDS) supplemented with 1X protease inhibitor (Roche cOmplete 11836153001). Cell lysates were vortexed and centrifuged, the supernatants were subjected to protein quantification by Bradford reagent (Bio-Rad 5000006) and sample preparation by sample buffer (10% glycerol, 50 mM Tris-HCl [pH 6.8], 2% SDS, 0.01% bromophenol blue and 8% β-mercaptoethanol). Protein samples were resolved by SDS-PAGE according to standard protocol and transferred onto 0.45μm nitrocellulose membranes (Bio-Rad 1620115) and blotted with the primary and secondary antibodies as described.

For Co-IP experiments, cells were lysed with RIPA buffer. The prepared protein extracts were precleared with protein A/G beads (Pierce, #20421) for 1 hours at 4 °C then incubated with anti-Flag M2 affinity gel for 2 hours for co-immunoprecipitation, followed by western blot analysis.

### Colony formation assays and WST-1 cell proliferation assays

Colony formation assays were done by seeding single cells in 6 or 12 well plates. Media was replenished every 3 days with indicated treatments. Colonies were fixed in 4% para-formaldehyde (PFA), followed by 0.5% crystal violet staining for 30 minutes at room temperature and rinsed with water. Quantification was performed using the ImageJ software plugin ColonyArea. Statistical significance was determined using unpaired, two-tailed Student’s t-test performed on intensity values from ColonyArea. For WST-1 cell proliferation assays (#11644807001, Roche), cells were seeded in 96 well plate for indicated days growth, and then were assayed according to the manufacturer’s instructions.

### RNA-sequencing

Cells from knockdown control or shWDR5-1 group were harvested with QIAzol Lysis Reagent (Qiagen) and homogenized using QIAshredder tubes (Qiagen). For each cell line, shRNA expression was induced with doxycycline (1 μg/mL) for 3 days and 3 biological replicates were harvested at different passages. RNA isolation was performed using miRNeasy with on-column DNase digestion. ERCC spike-in RNA was added in proportion to the number of cells obtained during cell counts. Library generation was performed using TruSeq stranded mRNA library prep kit (Illumina). Paired-end sequencing was performed using an Illumina HiSeq4000 sequencer, generating an average of 59 million reads per library. Reads were aligned to hg38 and gene counts to GENCODEv96 transcripts were obtained using STAR aligner v2.7.0 with default parameters. The hg38 and GENCODEv96 annotations were appended to include the ERCC sequences. DESeq2 was used to obtain differential gene expression, and HTSFilter was used to filter for expressed genes. Significant differences were identified using a BH adjusted p-value cut-off of 0.05. RNA-seq data have been deposited into the National Center for Biotechnology Information (NCBI) Gene Expression Omnibus database under GSE196666.

### RT-qPCR and barcode qPCR

Total RNA was extracted using the RNeasy Plus Mini Kit (Qiagen 74136) and reverse transcription was performed using High-Capacity cDNA Reverse Transcription kit (ThermoFisher 4385614). The resulting cDNA was diluted with water and Fast SYBR Green Master Mix (ThermoFisher 4385614) was used for real-time PCR. *GAPDH* was utilized as loading controls. Samples were run in quadruplicate and experiments were performed at least three times. Primer sequences are listed in **Table S5**. For barcode qPCR, barcode primers were designed to amplify only one barcode sequence among the 100 unique barcodes in the entire library. The primer set targeting the TRE element in pINDUCER10 was used for normalization. The full list of barcode qPCR primers used for detection of hairpin abundance is available in **Table S6**.

### Translation rate assay

Cells were starved of L-methionine for 30 minutes and subsequently incubated with 50 μM homopropargylglycine (HPG; Life Technologies #C10186) for 1 to 4 hours in treatment media. Cells were then trypsinized and fixed in 4% para-formaldehyde. A Click-IT kit (Life Technologies #C10269) used to label HPG. Labeled cells were analyzed using an Cytoflex flow cytometer. Translation rates were determined based on the slope of HPG incorporation over time. Significance determined using one sample *t* test to compare each treatment value to the hypothetical value 1.

### Chromatin Immunoprecipitation (ChIP)-qPCR

Cells growth in 15-cm dishes were washed with PBS and cross-linked with 1% formaldehyde in DMEM media for 10 minutes and quenched with 0.125 M glycine for 5 minutes. Cells were washed with cold PBS and scraped and pooled. Following washes, cell pellets were lysed in sonication buffer (20 mM Tris pH 8.0, 2 mM EDTA, 0.5 mM EGTA, 1X protease inhibitor, 0.5% SDS, 0.5 mM PMSF) at a concentration of 3 mL per 1×10^8^ cells for 10 minutes. Sonication was performed using the Qsonica Q800R sonicator (Qsonica) set to 70% amplitude, 15 seconds on and 45 seconds off for a total of 30 minutes on. Sonicated materials were pre-cleared with 50% protein A agarose (ThermoFisher 20421). Antibodies were added into pre-cleared material and rotated overnight at 4 °C. 50% protein A slurry was then added and tubes were rotated at 4 °C for 2 hours. In order to reverse crosslinks and purify DNA, NaCl was added to elute ChIP material and incubated overnight at 65 °C and then digested with proteinase K. Glycotube (ThermoFisher AM9515) was added as co-precipitant and phenol-chloroform isolation and ethanol precipitation was performed to isolate ChIP DNA. All sample DNA pellets were resuspended in 200 μL of water. 2 μL of DNA was used for each qPCR reaction, and reactions were performed in quadruplicate.

### 3D protein visualization

Protein crystal structure 2H14 (apo-WDR5) was downloaded from the Protein Data Bank and visualized using PyMol (The PyMOL Molecular Graphics System, Version 2.0 Schrdinger, LLC).

### Analysis of *in vitro* drug interaction

We employed coefficient of drug interaction to determine cytotoxicity. The coefficient of drug interaction (CDI) is calculated as follows: CDI=AB/(A×B). According to the colony formation intensity or translation rates of each group, AB is the ratio of the combination group to the control group; A or B is the ratio of the single agent group to the control group. Thus, CDI value<1, =1 or >1 indicates that the drugs are synergistic, additive or antagonistic, respectively. A CDI<0.7 indicates a significant synergistic effect (Otahal et al., 2020; Zhao et al., 2014).

### Statistical analysis

Comparisons between two groups were performed using an unpaired two-side Student’s *t* test. Graphs represent either group mean values ± SEM or individual values (as indicated in the figure legends). For animal experiments, each tumor graft was an independent sample. All experiments were reproduced at least three times.

## Acknowledgments

We would like to thank all members of Yan, Nguyen and Stern laboratories at Yale University for helpful discussions, Dr. Mei Zhong at Yale Stem Cell Center Genomics Core facility for helping with sample preparation for RNA-seq, Dr. Joan Massagué at Memorial Sloan Kettering Cancer Center for providing MDA-MB-231, LM2, and BoM cells, Dr. Yibin Kang at Princeton University for providing BrM2 cells, Dr. Narendra Wajapeyee at the University of Alabama Birmingham for helping with compiling the epigenetic gene list. Sequencing done at Yale Stem Cell Center Genomics Core facility was supported by the Connecticut Regenerative Medicine Research Fund and the Li Ka Shing Foundation. Figure panels 1A, 5A, and Figure 1-figure supplement 1A were created with BioRender.com.

## Author contributions

W.L.C., J.F.C., D.X.N., and Q.Y. designed the research. Q.Y. and D.X.N. conceived and oversaw the project. W.L.C. and J.F.C. performed most of the experiments. W.L.C. performed the bioinformatic analysis. W.L.C., J.F.C., L.H.C., A.A-E., M. Zhang and M. Zhao performed animal studies. W.L.C., J.F.C., H.C. and E.W. collected samples for RNA sequencing. W.L.C., J.F.C., S.J.K. and T.F.W. built the screening library. A.B. performed some in vitro assays. W.L. and Y.L. performed proteomic analysis. X.Y. and J.J provided degraders and helped with experimental design related to PROTAC. W.L.C., J.F.C., D.X.N., and Q.Y. analyzed the data. Y.D. contributed to analysis of data related to the KMT2 complexes. J.F.C., W.L.C., D.X.N., and Q.Y. wrote the paper.

## List of figure supplements

**Figure 1-figure supplement 1.**
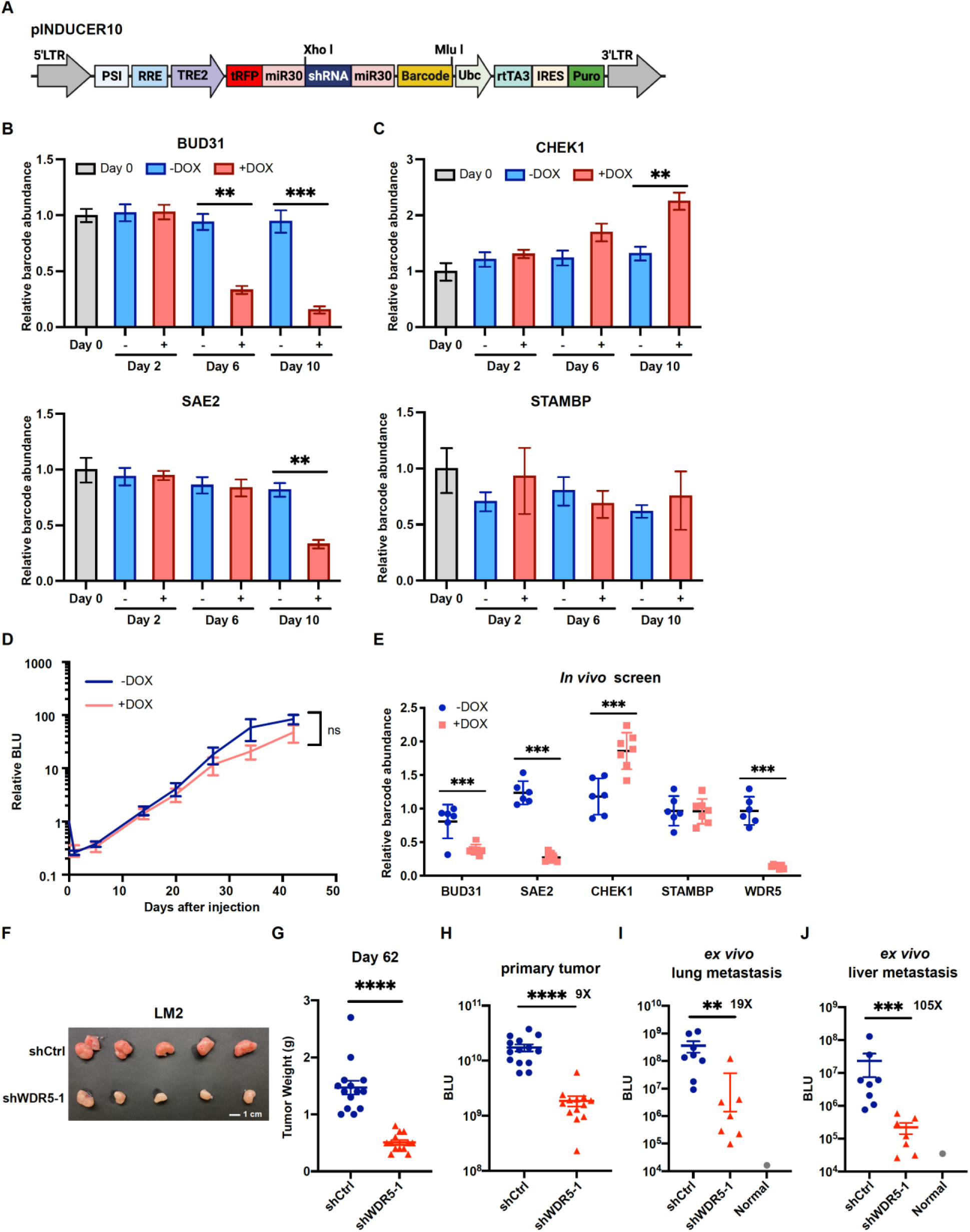
Positive controls and negative controls show expected phenotypes in both screening contexts. (A) Schematic representation of the pINDUCER10 plasmid. LTR, long terminal repeat; PSI, retroviral Ψ packaging element; RRE, Rev response element; TRE2, TRE2 promoter; Ubc, Ubiquitin C promoter; rtTA3, reverse tetracycline-controlled transactivator 3; IRES, internal ribosome entry site; Puro, puromycin resistance. (B-C) Relative abundance of cells stably expressing positive control shRNAs against *BUD31* and *SAE2* (B) and negative control shRNAs against *CHEK1* and *STAMBP* (C) after the indicated days of *in vitro* culture under control or doxycycline (1 μg/mL) treatment. Data normalized to abundance at the time of minipool mixture (Day 0). Representative data are shown from one minipool experiment (n=4, unpaired two-side Student’s *t* test). (D) Representative data of lung metastasis signal from mice injected with a minipool and fed with either doxycycline or regular chow at the indicated time point. ns, not significant. (E) Relative abundance of barcode for shRNA against *WDR5* and positive/negative controls in lung tissue from control and doxycycline-treated mice. (F) Representative image of primary tumor from mice injected into the 4^th^ mammary fat pad with LM2 cells harboring inducible control or shWDR5-1 at day 62. (G) Quantification of primary tumor weight from mice in (F) at day 62. Each dot represents one tumor. (H) Bioluminescence signals of the primary tumors from mice in (F) at day 61 post-injection (shCtrl: n=14; shWDR5: n=13). (I-J) Quantification of *ex vivo* bioluminescence signals of the lungs (I) and liver (J) from mice in (F) at day 62 post-injection. Each dot represents one animal (shCtrl: n=8; shWDR5: n=7). Significance determined using unpaired two-tailed Mann-Whitney test. *p<0.05; **p<0.001; ***p<0.001; ****p<0.0001.

**Figure 2-figure supplement 1.**
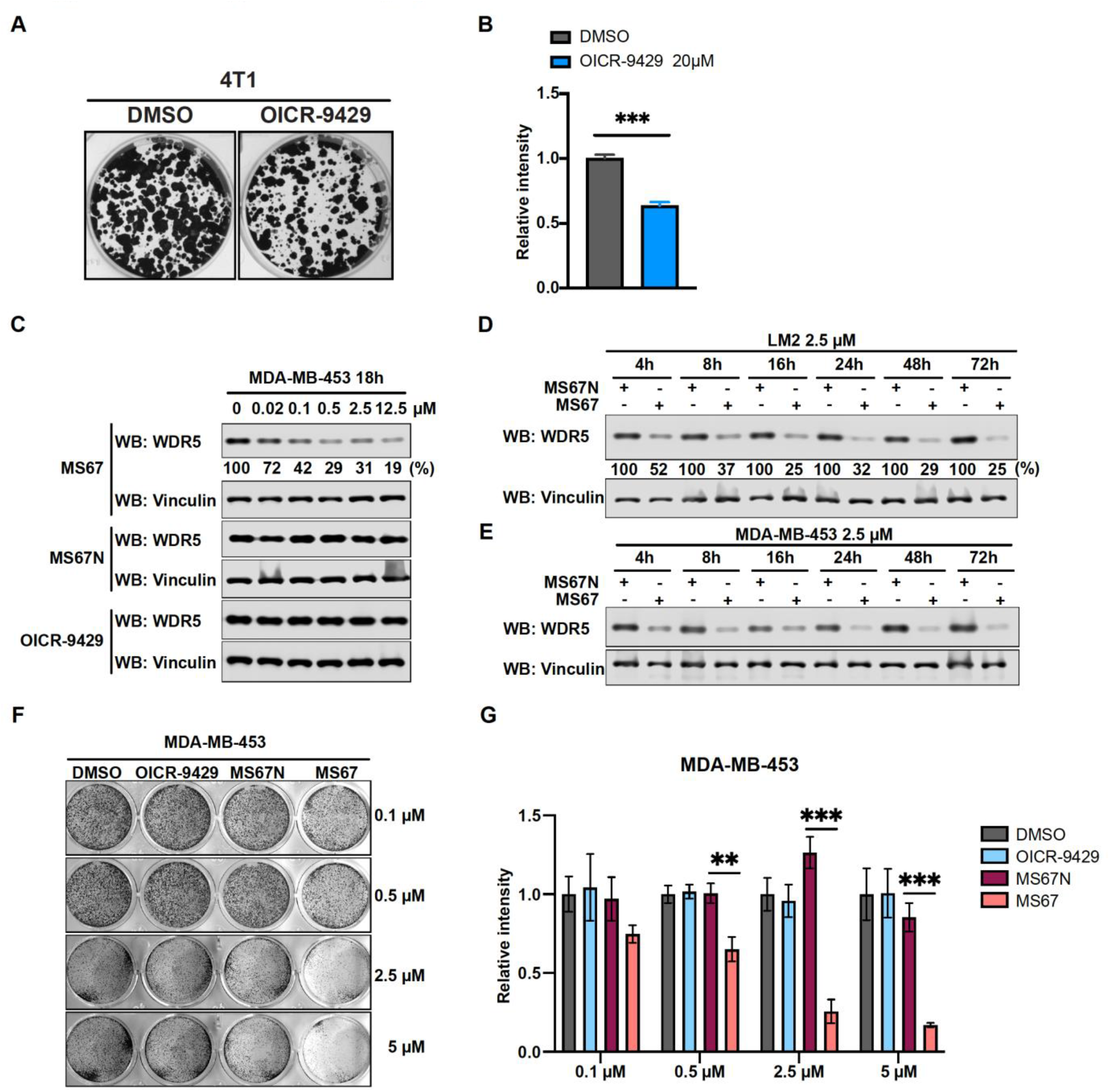
WDR5 inhibition and MS67-mediated WDR5 degradation significantly reduces breast cancer cell growth. (A-B) Colony formation assays of 4T1 after 9 days of either control or 20 μM OICR-9429 treatment for 9 days. Representative images (A) and quantification (B) are shown (n=3, unpaired two-side Student’s *t* test). (C) Western blot analysis of WDR5 in MDA-MB-453 treated with MS67, MS67N, or OICR-9429 at indicated concentrations for 18 hours. Band intensities were quantified by image J. (D-E) Western blot analysis of WDR5 in LM2 (D) or MDA-MB-453 (E) treated with 2.5 μM MS67 or MS67N for the indicated durations. Band intensities were quantified by image J and normalized by the those of vinculin control. (F-G) Colony formation assays of MDA-MB-453 after 9 days of treatment with control, OICR-9429, MS67N, or MS67 at the indicated concentrations. Representative images (F) and quantification (G) are shown (n=3, unpaired two-side Student’s *t* test). *p<0.05; **p<0.001; ***p<0.001; ****p<0.0001. For gel source data, see Figure 2- figure supplement 1-source data 1-3.

**Figure 3-figure supplement 1.**
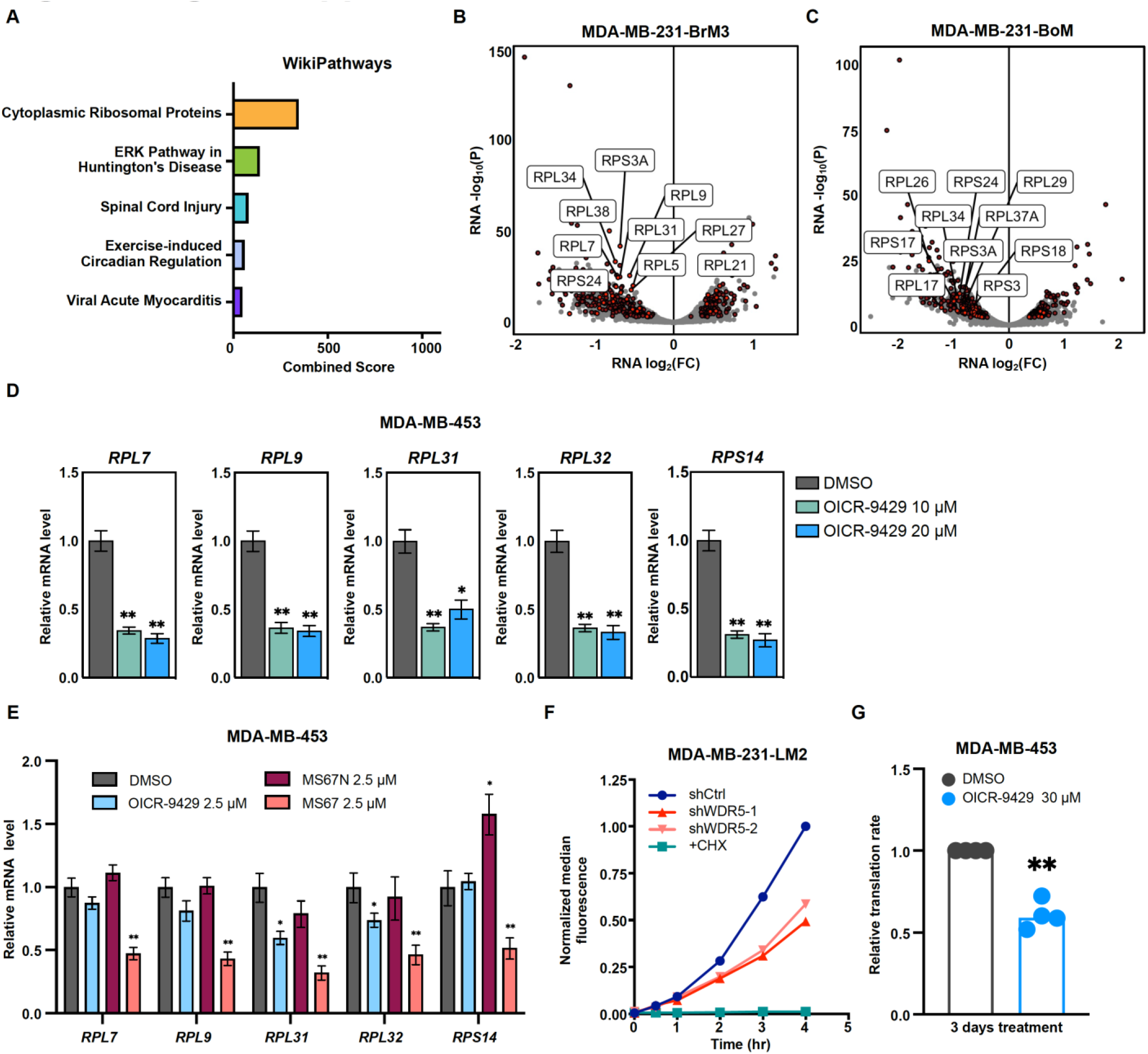
WDR5 targeting decreases ribosomal protein gene expression and global translation rates. (A) Gene ontology results using the up-regulated gene set shared by three MDA-MB-231 organotropic sublines analyzed with Enrichr. (B-C) Volcano plot of genes differentially expressed after WDR5 knock-down in BrM3 (B) and BoM (C). Shared DEGs across all lines highlighted in dark red and RPs (RPL and RPS) highlighted in light red. The top ten differentially expressed RPs are labelled. (D) RT-qPCR validation of selected DEGs in MDA-MB-453 cells after control or OICR-9429 treatments at the indicated concentrations for 3 days. (E) RT-qPCR validation of selected DEGs in MDA-MB-453 cells after DMSO, 2.5 μM OICR-9429, MS67N, or MS67 treatment for 3 days. Significance determined by comparing each treatment to DMSO control (n=4, unpaired two-side Student’s *t* test). (F) Representative translation efficiency overtime using indicated LM2 cell lines with continuous treatment of doxycycline (1 μg/mL). 100 μg/mL cycloheximide (CHX) was used as a control to completely block *de novo* protein translation. (G) Normalized translation rates of MDA-MB-453 cells following 3 days of control or 30 μM OICR-9429 treatment (n=4, one sample t-test). Significance determined using one sample t-test. *p<0.05; **p<0.001; ***p<0.001; ****p<0.0001.

**Figure 4-figure supplement 1.**
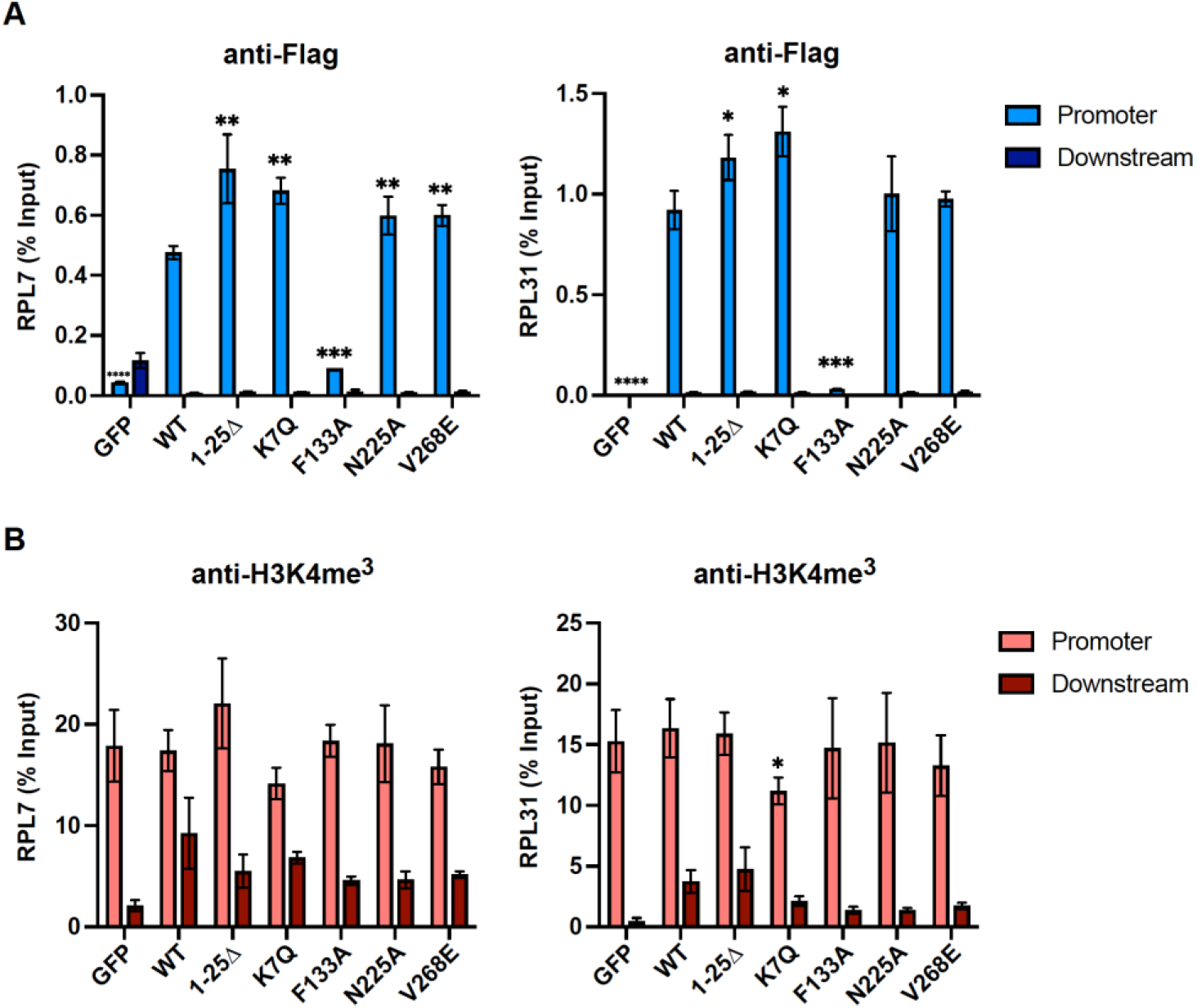
WDR5 recruitment to ribosomal protein gene promoters is not sufficient for gene activation. (A-B) ChIP-qPCR data of the indicated over-expression condition using primers for either the promoter or 800 bp downstream control at the RPL7 (left) or RPL31 (right) gene locus. ChIP was performed using the anti-Flag to pull down WDR5 (A) and H3K4me3 (B). Significance determined by comparing promoter enrichment of WT to other over-expressing conditions using unpaired two-tailed Student’s t-test.

**Figure 6-figure supplement 1.**
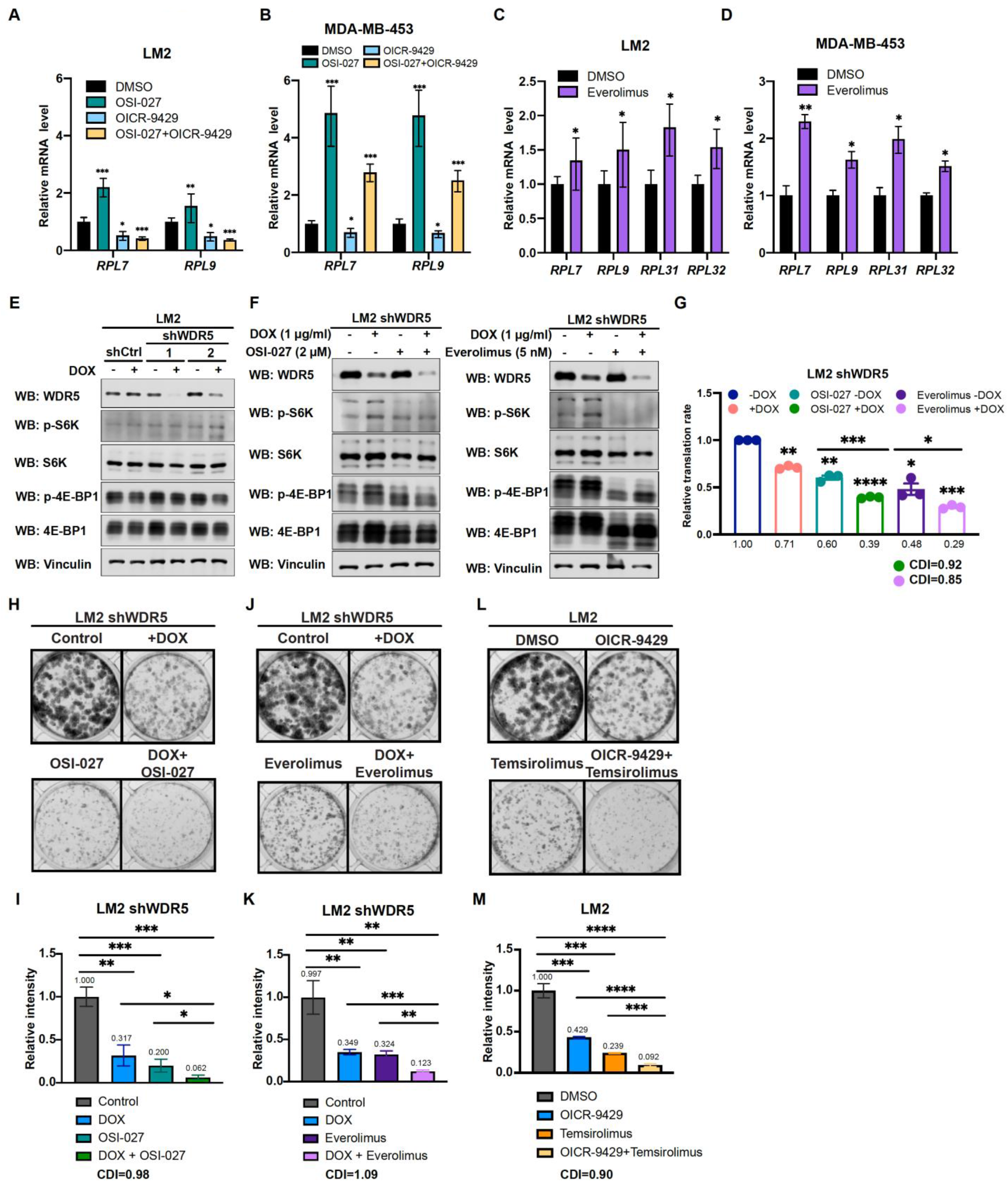
Inhibition of WDR5 and mTOR cooperatively reduces translation and TNBC growth. (A) RT-qPCR of selected DEGs in LM2 after DMSO, 2 μM OSI-027, 20 μM OICR-9429, or combined treatment for 3 days. (B) RT-qPCR of selected DEGs in MDA-MB-453 after DMSO, 0.5 μM OSI-027, 30 μM OICR-9429, or combined treatment for 3 days. (C) RT-qPCR of selected DEGs in LM2 after DMSO or 5 nM everolimus treatment for 3 days. (D) RT-qPCR of selected DEGs in MDA-MB-453 after DMSO or 1 nM everolimus treatment for 3 days. Significance determined by comparing each treatment to DMSO control (n=4, unpaired two-side Student’s *t* test). (E) Western blot analysis of the indicated proteins in LM2 shCtrl, shWDR5-1, and shWDR5-2 with or without hairpin induction by doxycycline (1 μg/mL) for 3 days. (F) Western blot analysis of indicated proteins in LM2 shWDR5-1 with or without doxycycline (1 μg/mL) induction in combination with 3 days of control, 2 μM OSI-027, or 5 nM everolimus treatment. (G) Normalized translational rates of LM2 shWDR5 cells from (A) following 3 days of control, 2 μM OSI-027, or 5 nM everolimus treatment (n=3, one sample t-test). (H-I) Colony formation assay of LM2 shWDR5 with or without hairpin induction by doxycycline in combination with control or 2 μM OSI-027 treatment for 8 days. Representative images (H) and quantification (I) are shown. (J-K) Colony formation assay of LM2 shWDR5 cells with or without hairpin induction by doxycycline in combination with control or 5 nM everolimus treatment for 8 days. Representative images (J) and quantification (K) are shown. (L-M) Colony formation assay of LM2 with or without 20 μM OICR-9429 in combination with control or 2.5 μM temsirolimus treatment for 8 days. Representative images (M) and quantification (L) are shown (n=3, unpaired two-side Student’s *t* test). *p<0.05; **p<0.001; ***p<0.001; ****p<0.0001. Calculation of coefficients of drug interaction (CDI) is described in materials and methods section. Significant synergy is labeled with (#). For gel source data, see Figure 6- figure supplement 1-source data 1-2.

## List of source data

**Figure 1-source data 1:** Original western blots for Figure 1D

**Figure 2-source data 1:** Original western blots for Figure 2A

**Figure 2-source data 2:** Original western blots for Figure 2H

**Figure 2-figure supplement 1-source data 1:** Original western blots for Figure 2-figure supplement 1C

**Figure 2- figure supplement 1-source data 2:** Original western blots for Figure 2-figure supplement 1D

**Figure 2- figure supplement 1-source data 3:** Original western blots for Figure 2-figure supplement 1E

**Figure 4-source data 1:** Original western blots for Figure 4C

**Figure 4-source data 2:** Original western blots for Figure 4D

**Figure 5-source data 1:** Original western blots for Figure 5B

**Figure 6-source data 1:** Original western blots for Figure 6A

**Figure 6-source data 2:** Original western blots for Figure 6B

**Figure 6- figure supplement 1-source data 1:** Original western blots for Figure 2-figure supplement 1E

**Figure 6- figure supplement 1-source data 2:** Original western blots for Figure 2-figure supplement 1F

**Figure 7-source data 1:** Original western blots for Figure 7A

**Figure 7-source data 2:** Original western blots for Figure 7B

